# Phosphatidylinositol 4-kinase III alpha governs cytoskeletal organization for invasiveness of liver cancer cells

**DOI:** 10.1101/2023.05.22.541742

**Authors:** Cong Si Tran, Julia Kersten, Marco Breinig, Jingyi Yan, Tanja Poth, Ombretta Colasanti, Tobias Riedl, Suzanne Faure-Dupuy, Stefan Diehl, Lieven Verhoye, Teng- Feng Li, Marit Lingemann, Philipp Schult, Gustaf Ahlén, Lars Frelin, Florian Kühnel, Kai Breuhahn, Florian W. R. Vondran, Philip Meuleman, Mathias Heikenwälder, Peter Schirmacher, Matti Sällberg, Ralf Bartenschlager, Vibor Laketa, Darjus Felix Tschaharganeh, Volker Lohmann

**Author notes:** Corresponding author: Prof. Dr. Volker Lohmann, Heidelberg University, Medical Faculty Heidelberg, Department of Infectious Diseases, Molecular Virology, Section Virus-Host-Interactions, Center for Integrative Infectious Disease Research, Heidelberg, Im Neuenheimer Feld 344, 69120, Germany.

## Abstract

**Background and Aims:** High expression of phosphatidylinositol 4-kinase III alpha (PI4KIIIα) correlates with poor survival rates in patients with hepatocellular carcinoma (HCC). In addition, Hepatitis C virus (HCV) infections activate PI4KIIIα and contribute to HCC progression. We aimed at mechanistically understanding the impact of PI4KIIIα on the progression of liver cancer and the potential contribution of HCV in this process.

**Methods:** Several hepatic cell culture and mouse models were used to study functional importance of PI4KIIIα on liver pathogenesis. Antibody arrays, gene silencing and PI4KIIIα specific inhibitor were applied to identify the involved signaling pathways. The contribution of HCV was examined by using HCV infection or overexpression of its nonstructural protein.

**Results:** High PI4KIIIα expression and/or activity induced cytoskeletal rearrangements via increased-phosphorylation of paxillin and cofilin. This led to morphological alterations and higher migratory and invasive properties of liver cancer cells. We further identified the liver specific lipid kinase phosphatidylinositol 3-kinase C2 domain-containing subunit gamma (PIK3C2γ) working downstream of PI4KIIIα in regulation of the cytoskeleton. PIK3C2γ generates plasma membrane (PM) phosphatidylinositol 3,4-bisphosphate [PI(3,4)P2]- enriched, invadopodia-like structures which regulate cytoskeletal reorganization by promoting Akt2 phosphorylation.

**Conclusions:** PI4KIIIα regulates cytoskeleton organization via PIK3C2γ/Akt2/paxillin-cofilin to favor migration and invasion of liver cancer cells. These findings provide mechanistic insight into the contribution of PI4KIIIα and HCV to progression of liver cancer and identify promising targets for therapeutic intervention.

**IMPACT AND IMPLICATIONS:** Understanding mechanistically how high PI4KIIIα expression are associated with poor clinical outcomes of liver cancer is important to develop pharmaceutical interventions. Our study sheds light on the importance of the two lipid kinases PI4KIIIα and PIK3C2γ as well as the contribution of HCV on liver cancer progression, unraveling the signaling pathway governing this process. This preclinical study contributes to better understanding the complex connection of phospholipids, cytoskeleton and liver cancer and suggests strategies to improve therapeutic outcomes by targeting important signaling molecules.

**Graphical abstract:** 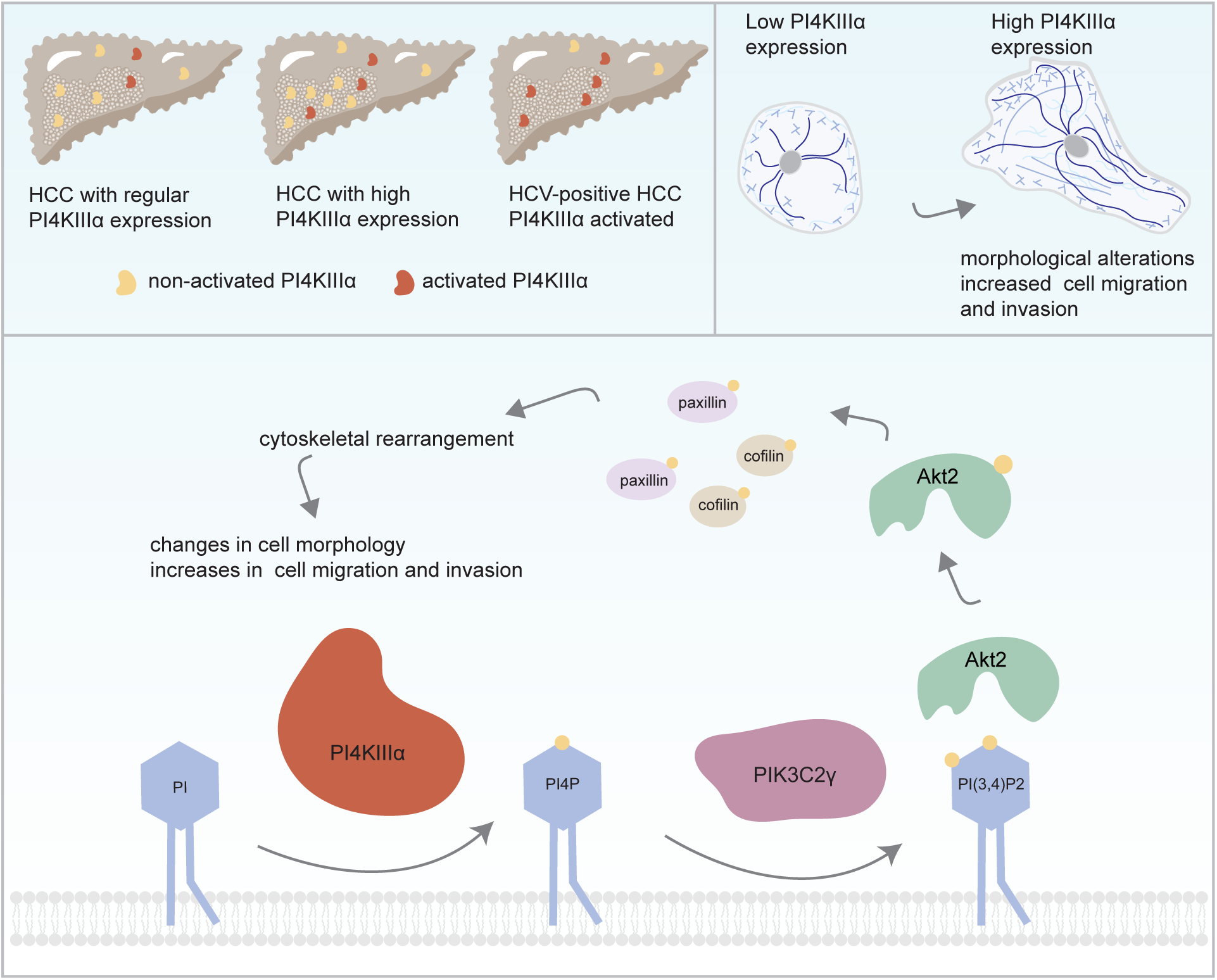

## INTRODUCTION

With an estimated incidence of 900.000 cases, accounting for more than 830.000 deaths worldwide in 2020 [1], liver cancer remains a public health challenge and is predicted to rise exponentially in the coming years. HCC is the most common primary liver malignancy and the fourth leading cause of cancer-related death globally [1].

Malignant transformation of normal cells to cancer cells is a multi-step process in which changes in cell morphology and motility are frequently observed. Cytoskeleton reorganization plays an active role in inducing these changes which requires concerted actions of a plethora of cytoskeletal proteins [2]. Coordination of cytoskeletal dynamics critically involves participation of phosphoinositides [3], and thus these signaling entities are in addition important in cancer progression [4]. The interconversion of the seven phosphoinositides and phosphatidylinositol (PI) by different phospholipid kinases, phosphatases and phosphoinositide modifying enzymes is the signature of the phosphoinositide signaling system [4].

Among the phosphoinositide lipids, phosphatidylinositol 4-phosphate (PI4P) is the most abundant monophosphorylated derivative of PI [5]. The conversion to PI4P from PI requires a phosphatidylinositol 4-kinase (PI4K) [6]. Human PI4Ks comprise 4 members with their localization defining different spatial distributions of cellular PI4P pools and thus divergence of their cellular functions [6]. PI4KIIIα is the largest member of the PI4Ks family, resides in the endoplasmic reticulum and transiently shuttles to the PM, mediated by its N-terminus [7, 8]. Mutations in the catalytic domain, e.g. D1957A, located at the very C-terminus [7], render these variants catalytically inactive. Notably, only trace amounts of PI4KIIIα are present in normal liver, but its expression is highly upregulated and associated with hepatic dedifferentiation and poor prognosis in HCCs [9]. Although the functional importance of this lipid kinase in HCV replication has been intensely studied and several PI4KIIIα inhibitors have been developed for antiviral strategies [8], it remains unclear if PI4KIIIα activation contributes to virus-induced cancer development.

In this study, we aimed to decipher PI4KIIIα effects related to progression of liver cancer. We found that increased abundance and activity of PI4KIIIα resulted in elevated PI(3,4)P2 levels in invadopodia-like structures at the PM, mediated by the liver specific lipid kinase PIK3C2γ. In turn, the activation of Akt2 promoted increased phosphorylation of paxillin and cofilin, correlating with migratory and invasive properties of liver cancer cells and respective morphological changes. HCV infection further promoted these phenotypes by activating PI4KIIIα. Hereby we identified a molecular pathway based on high PI4KIIIα expression and activity promoting liver carcinogenesis.

## MATERIALS AND METHODS

### Cell lines and culture conditions

All eukaryotic cells were cultured in Dulbecco’s modified Eagle’s medium (DMEM; Gibco) containing 10% FCS (Seromed), 1% penicillin/streptomycin (Gibco), 2 mM L-glutamine (Gibco) and 1% non-essential amino acids (Gibco) or HCM Hepatocyte Culture Medium BulletKit (for HHT4) at 37 °C in a constant humid atmosphere containing 5% CO2. Cells were regularly tested for Mycoplasma contamination using a commercially available system (MycoAlert Mycoplasma Detection kit, Lonza).

### Generation of stable knockdown or overexpression cell lines via lentiviral transduction

One day prior to transfection, 6×10^6^ 293T cells were seeded per 10 cm-diameter dish in DMEM complete. 30 min before transfection, medium was replaced by serum-free DMEM. For transfection, 5.15 µg packaging plasmid (pCMV-Gag-Pol), 1.72 µg of the VSV envelope glycoprotein expression vector (pMD2-VSVG) and 5.15 µg of transfer vector encoding either the respective shRNAmir (and puromycin resistance (pAPM)) or the respective shRNAmir-escape variants (and blasticidin resistance (pWPI) for expression of PI4KIIIα, paxillin, cofilin and Akt2 variants) were mixed in 360 µL Opti-MEM. In a separate tube 36 µL polyethylenimine (PEI) was mixed with 360 µL Opti-MEM. The PEI mixture was then added dropwise to the plasmid mixture, followed by vortexing for 10 seconds and 20 min incubation at room temperature (RT). The transfection mixture was then added dropwise to different area of the dishes and was distributed evenly by gently rocking the culture vessel back-and-forth and from side-to-side. 6 h post-transfection medium was replaced with DMEM complete, followed by 3 days incubation. Lentivirus-containing-supernatant was harvested and filtered through a 0.45 µm filter and was used to transduce target cells. Cells were then subjected to selection with medium containing appropriate antibiotics (2.5 µg/mL puromycin and/or 10 µg/mL blasticidin).

### Cell invasion assay

Cell invasion was evaluated out using BioCoat Matrigel Invasion Chamber (Corning) according to manufacturer’s instructions. In brief, prior to cell seeding, Matrigel inserts in 24-well plate were rehydrated with serum-free DMEM for 2 h in humidified tissue culture incubator (37 °C, 5% CO2). In upper chamber, cells were seeded at the density of 1.25×10^4^ per well (0.5 mL) in serum-free DMEM. In lower chamber under the insert, 0.75 mL complete DMEM was added to serve as a chemoattractant. Cells were incubated for 20 h in humidified tissue culture incubator. Invading cells were stained by 0.5% Crystal violet and counted.

### Quantification and statistical analysis

Statistical analyses were performed using two-tailed, unpaired *t*-test with Welch’s correction in GraphPad Prism v6 and v7. Data are presented as means ± standard deviations (SD). The asterisks in the figures indicate significant differences: not significant (ns, p > 0.05), * (p ≤ 0.05), ** (p ≤ 0.01), *** (p ≤ 0.001) and **** (p ≤ 0.0001). Immunoblot analyses and confocal immunofluorescence were processed and quantified using Fiji. Images from cell culture microscopy were processed and analyzed using Apeer, ilastik and Fiji. ImageScope and Fiji were used to process and analyze immunohistochemistry data. Arrangement of figures was carried out using Inkscape and Adobe Illustrator.

## RESULTS

### PI4KIIIα expression and/or activity links to cell morphology

Human liver cancer cell lines show higher PI4KIIIα expression than PHHs [9, 10]. To understand the functional importance of this lipid kinase for liver pathogenesis, we firstly silenced *PI4KA* in Huh7-Lunet cells (Fig. 1A-B). Although cell proliferation remained unchanged (Fig. S1A), *PI4KA* silencing led to drastic alterations in cell morphology (Fig. 1C). HCC-derived Huh7-Lunet cells evenly spread and exhibited hexagonal to rounded shape, a phenotype designated as “cancer cell morphology” throughout the manuscript. In contrast, *PI4KA* knockdown cells showed diverse compact shapes. Both quantification with manual classification (Fig. 1D) and automated classification using deep learning confirmed a significant drop of cancer cell morphology in the cells lacking PI4KIIIα (Fig. 1E). The same morphological alterations due to differences in PI4KIIIα expression were observed in two additional human liver cancer cell lines, Huh7.5 and Hep3B (Fig. S1B-E). PI4KA-F1, a selective inhibitor of PI4KIIIα [8], induced similar changes in the morphology of Huh7-Lunet (Fig. 1F-G), Huh7.5 (Fig. S1F-G) and Hep3B (Fig. S1H-I) cells, in a time and dose dependent manner (Fig. 1G and S1G). Expression of wild-type PI4KIIIα (*PI4KA* wt), but not catalytically inactive PI4KIIIα (*PI4KA* mut) in *PI4KA* knock-down cells reverted the morphological changes (Fig. 1H-I). Taken together, these data suggested a strong association of PI4KIIIα expression and activity with tumorous cytomorphology.

**Figure 1.**
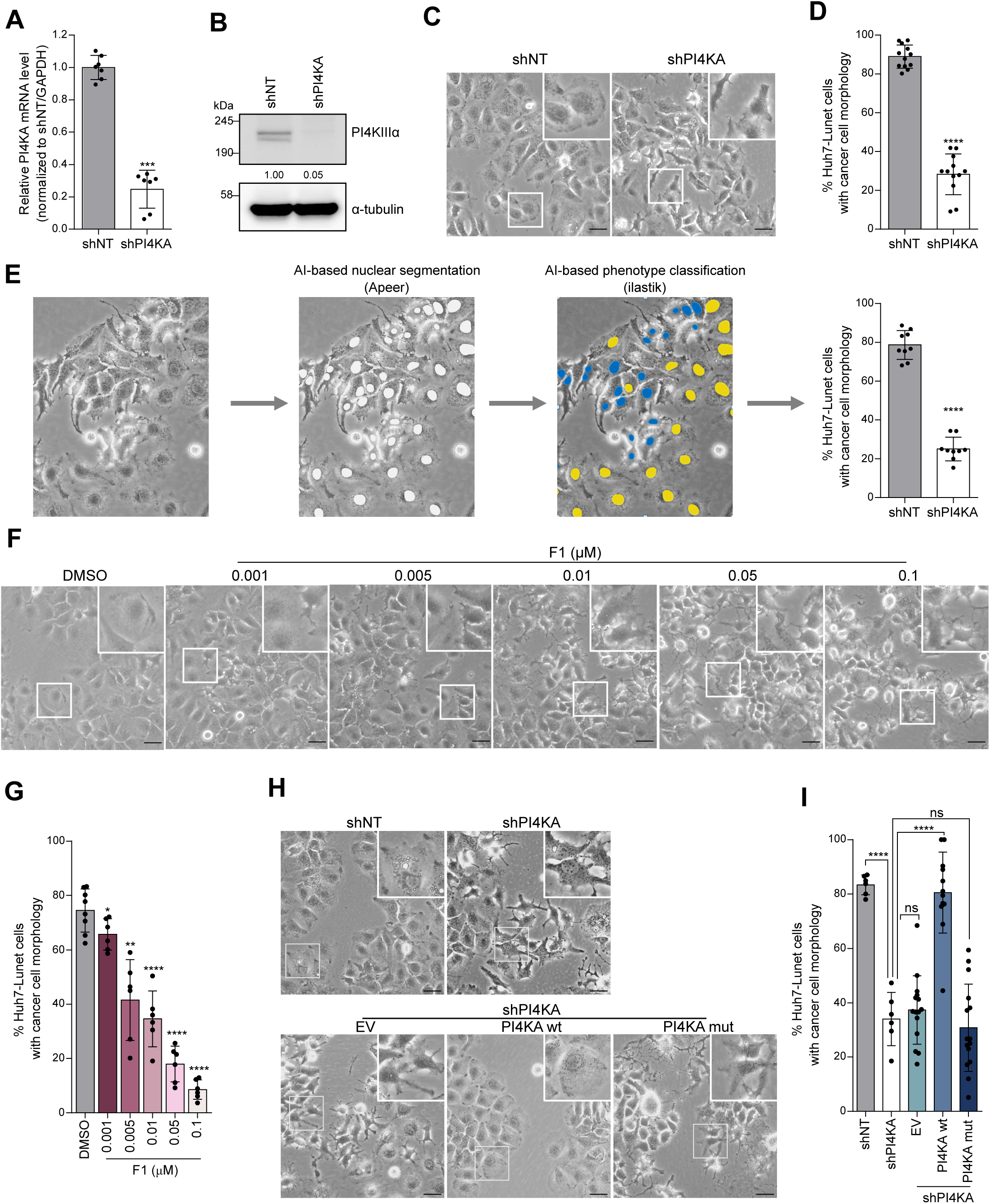
PI4KIIIα links to cell morphology. (A-D) Knockdown efficiency of shPI4KA in Huh7-Lunet cells assessed by RT-qPCR (A) or WB (B). Numbers below each lane represent relative expression levels compared to shNT. Cells were then imaged for cell morphology (C) and cancer cell morphology was quantified by manual classification (D). (E) Schematic workflow for automated classification of cell morphology by a machine learning approach. (F-I) Visualization of cell morphology (F, H) and quantification of cancer cell morphology (G, I) in Huh7-Lunet cells either treated with DMSO or PI4KA-F1 for 48 h (F, G) or expressing PI4KIIIα wt or an inactive mutant (H, I).

### PI4KIIIα expression and activity favor cell migration and invasion

We next asked if PI4KIIIα was involved in cell migration and cell invasion. We found that both *PI4KA* silencing (Fig. 2A-B) and inhibitor PI4KA-F1 treatment (Fig. S2A-B) markedly reduced cell migration. In accordance with the cell migration assay, we observed a reduced invasion in Huh7-Lunet cells expressing shPI4KA (Fig. 2C-D) or treated with PI4KA-F1 (Fig. 2E). This observation was further confirmed upon silencing of *PI4KA* in mouse hepatoma cells Hep55.1C (Fig. S2C-D). Furthermore, only expression of *PI4KA* wt restored the invasiveness of shPI4KA cells (Fig. 2F). In addition, we found that E-cadherin expression was consistently higher in *PI4KA*-silenced cells (Fig. S2E), indicating a reversion of the epithelial–mesenchymal transition process, concurrent with reduced invasiveness (Fig. 2C-D).

**Figure 2.**
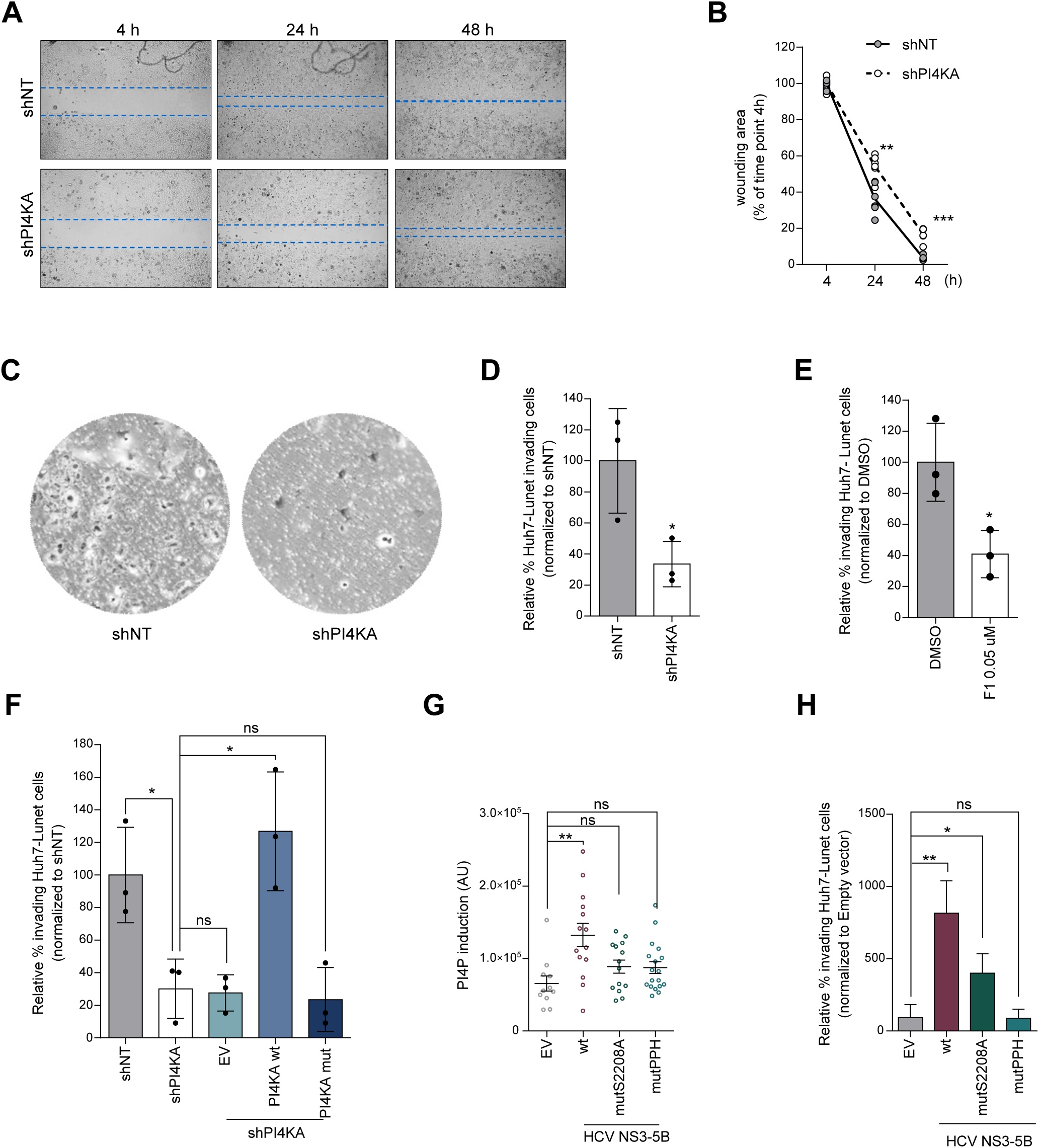
PI4KIIIα facilitates liver cancer cell motility. (A-B) Migration assay was performed with Huh7-Lunet-shNT and -shPI4KA cells (A) and wounding areas were analyzed (B). (C-D) Invasion assay was performed with Huh7-Lunet-shNT and -shPI4KA. Invading cells were stained (C) and relative cell invasion rate was evaluated (D). (E-F) Relative invasion rate of Huh7-Lunet cells treated with DMSO or PI4KA-F1 (E) and Huh7-Lunet-shPI4KA cells expressing PI4KIIIα variants (F). (G-H) Levels of PI4P (in at least 10 cells) (G) and relative percentage of invading cells of Huh7-Lunet expressing the indicated HCV NS3-5B variants (H).

To assess the impact of HCV-activated PI4KIIIα on invasiveness, we expressed parts of the HCV polyprotein, encompassing nonstructural proteins NS3-5B upon transduction of Huh7-Lunet cells with adenoviral vectors. We observed in cells expressing HCV NS3-5B wt a significant induction of PI4P (indicating PI4KIIIα activation) and in turn an increase in cell invasion which was lower in case of mutant S2208A or completely abrogated in cells expressing the mutant PPH (Fig. 2G-H). A similar effect was also observed in HHT4 cells, a non-transformed, immortalized hepatocyte cell line (Fig. S2F).

In conclusion, these data suggested a significant contribution of both expression and activity of PI4KIIIα to cell motility, which may contribute to invasion and metastatic spread.

### The cytoskeletal network is regulated by PI4KIIIα via enhanced phosphorylation of paxillin and cofilin

Considering the key role of the cytoskeleton in all morphogenesis processes as well as in cell migration and invasion [2], we speculated that PI4KIIIα promotes reorganization of the cytoskeletal network. Indeed, actin fibers, microtubules and intermediate filaments were evenly distributed throughout the cell body of the shNT cells but highly condensed in the *PI4KA* knockdown cells (Fig. 3A-B and S3A).

**Figure 3.**
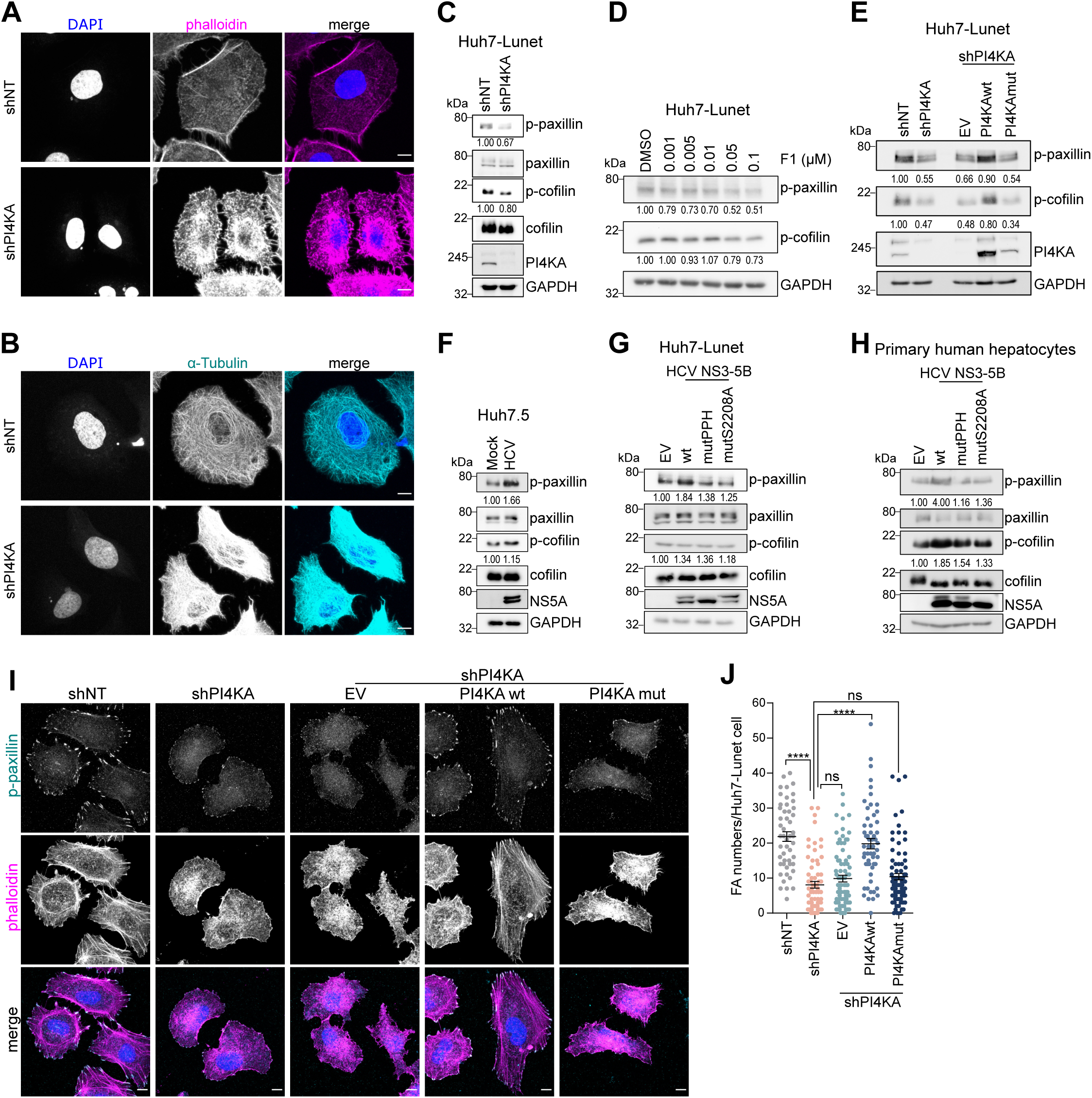
The cytoskeletal network is regulated by PI4KIIIα via an enhanced phosphorylation of paxillin and cofilin. (A-B) F-actin (A) and tubulin (B) in Huh7-Lunet-shNT and –shPI4KA. (C-H) Immunoblotting of p-paxillin and p-cofilin in Huh7-Lunet cells, either expressing shNT/shPI4KA (C) or treated with PI4KA-F1 (D), Huh7-Lunet-shPI4KA cells (E), Huh7.5 cells infected with HCVcc Jc1 at MOI=0.5 for 4 days (F), Huh7-Lunet cells (G) or PHH (H) expressing the indicated HCV NS3-5B variants. Numbers below each lane represent relative expression levels compared to the respective control shNT, DMSO, mock infection or EV. (I-J) FAs indicated by p-paxillin staining (I) and quantification of FAs numbers per cell (in at least 50 cells) (J) in Huh7-Lunet cells expressing PI4KIIIα variants.

In search of a possible link between PI4KIIIα and the cytoskeleton reorganization, we performed a cytoskeleton phospho-antibody array and identified p-paxillin Y31 and cofilin, as candidate proteins regulated by PI4KIIIα activity (Fig. S3B and Table S1). Another screening in HEK293 cells also revealed similar changes of these two proteins by PI4KIIIα abundance [11]. Moreover, paxillin and cofilin play key roles in structuring cytoskeletal networks and their phosphorylation is required for cancer cell migration, invasion and chemotherapy resistance [12-15]. We therefore further investigated the regulation of paxillin and cofilin by PI4KIIIα in liver cells. PI4KA silencing led to a significant reduction in the levels of p-paxillin (Y31) and a slight reduction in p-cofilin (S3) abundance, while the total expression remained unchanged (Fig. 3C). These effects were consistently phenocopied in a panel of liver-cancer cell lines, including Huh7, Huh7.5, Hep3B and HepG2 (Fig. S3C-F). PI4KA-F1 treatment also led to a dose-dependent reduction of p-paxillin and p-cofilin (Fig. 3D). Moreover, p-paxillin and p-cofilin levels were restored by the re-expression of *PI4KA* wt in *PI4KA* knockdown cells (Fig. 3E). Paxillin and cofilin mutants harboring phosphomimetic mutations Y31D or S3D, respectively, significantly reverted the effect of shPI4KA on cell morphology (Fig. S3G-H). Taken together, our data suggest that PI4KIIIα abundance and activity are required for phosphorylation of paxillin and cofilin in liver cancer cells which are important steps mediating the impact of increased PI4P levels to cytoskeletal changes.

We next examined the effects of HCV on paxillin and cofilin phosphorylation. We confirmed an induction of PI4P in HCV-infected cells (Fig. S3I) and in turn observed an increase in levels of p-paxillin and to a lesser extent of p-cofilin (Fig. 3F). In addition, in the replication-independent NS3-5B expression system, levels of increased phospho-paxillin (Fig. 3G) correlated with the induction of PI4P in these cells (Fig. S2A) which was further confirmed in PHHs (Fig. 3H), indicating that HCV-induced PI4P mediated changes can also be observed at physiological PI4KIIIα concentrations.

Paxillin is a key component of focal adhesions (FAs), which contribute significantly to regulation of cell shape and cell motility. Silencing *PI4KA* reduced significantly FA numbers (Fig. 3I-J) which could only be reverted to the levels measured in the shNT sample by reintroducing PI4KIIIα wt (Fig. 3I-J). In contrast, HCV infection led to approximately 3-fold increase in FA numbers (Fig. S3J-K). These data highlight the positive correlation between PI4KIIIα activity and FA numbers and its contribution to cell motility via phosphorylation of paxillin.

### PI4KIIIα expression/activity and phosphorylation levels of paxillin and cofilin correlate *in vivo*

In order to validate the impact of PI4KIIIα on paxillin and cofilin phosphorylation *in vivo*, we exogenously expressed PI4KIIIα or HCV NS3-5B in the liver of mice. We used a Sleeping Beauty transposon vector, capable of integrating the sequence of interest into the genome of transfected cells [16], or a plasmid harboring a scaffold/matrix attachment region (pS/MARt) [17], allowing persistent gene expression. The constructs were tested for expression in cell culture (Fig. S4A) before delivery via hydrodynamic tail-vein injection. Mouse livers expressing PI4KIIIα wt or HCV NS3-5B wt contained significantly higher levels of p-paxillin compared to empty vector controls or inactive mutants, whereas p-cofilin amounts were only slightly increased (Fig. 4A-B and S4B). PI4KIIIα wt and HCV wt–expressing mouse livers also showed higher grades of steatosis and hepatocellular hypertrophy (Fig. S4C-D). We next analyzed p-paxilin levels in human-liver chimeric mice which are susceptible for HCV infection [18]. Expression of human albumin and HCV NS5A confirmed a successful HCV infection in the human hepatocytes engrafted in the mice (Figure 4C-D). Moreover, p-paxillin was much more abundant in HCV-infected samples (Fig. 4C-D and S4E-F). These data establish the *in vivo* relevance of our *in vitro* findings that PI4KIIIα and HCV-activated PI4KIIIα regulate the organization of the cytoskeleton by increasing p-paxillin and p-cofilin levels.

**Figure 4.**
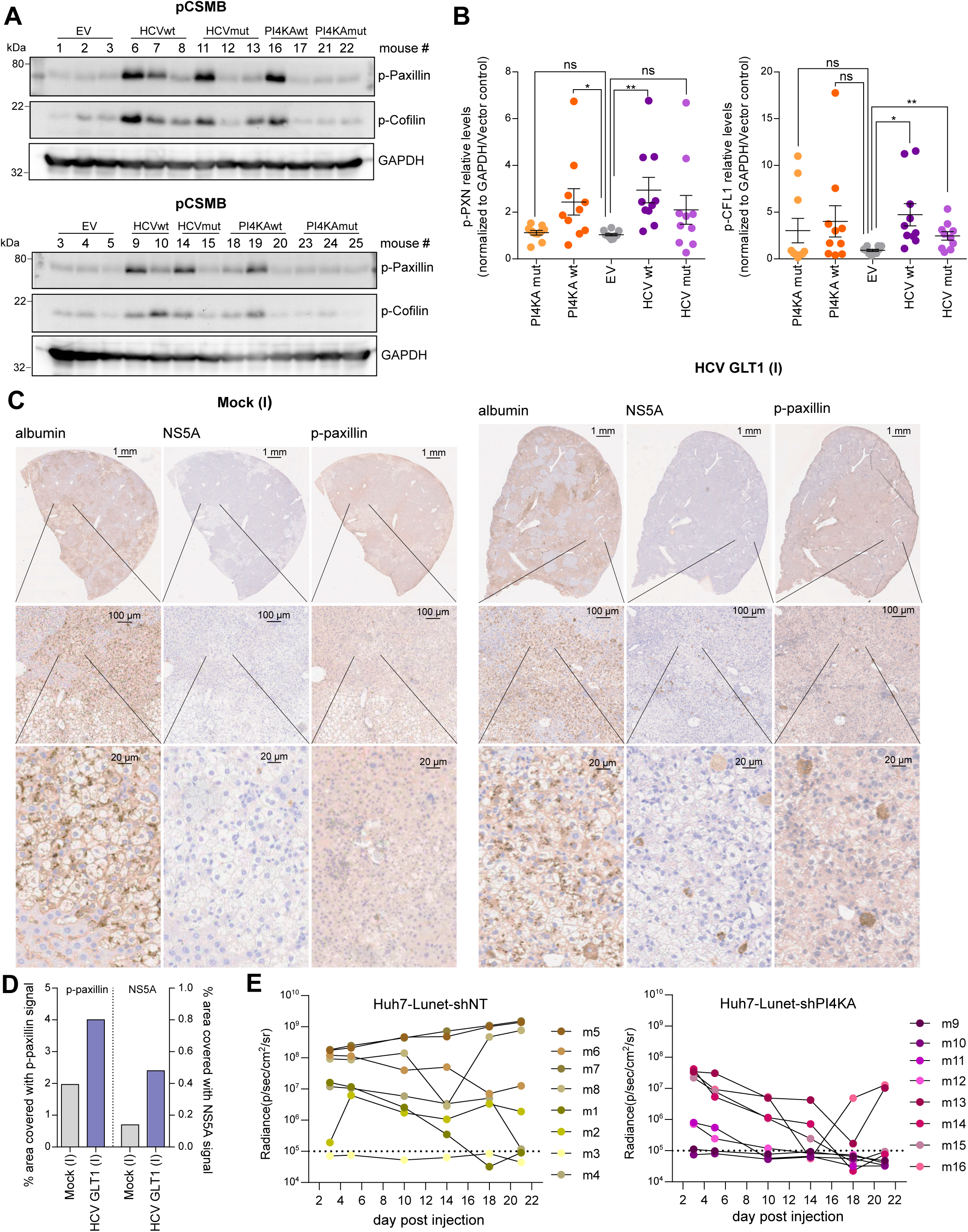
PI4KIIIα expression/activity and phosphorylation levels of paxillin and cofilin correlate *in vivo*. (A-B) Immunoblotting of p-paxillin and p-cofilin in liver of mice expressing the indicated PI4KIIIα/HCV NS3-5B variant (via hydrodynamic tail vein injection with pCSMB vectors) (A, also in S4B) and the relative levels were quantified (B). 10 mice for each treatment group. (C) Staining of human albumin, HCV NS5A and p-paxillin in livers of human-liver chimeric mice infected with HCV patient serum (gt 1b, isolate GLT1). (D) Fraction of the area of the histology sections covered with p-paxillin or NS5A signals in mock and HCV GLT1 infected samples was automatically determined using the random forest model trained in ilastik. (E) Huh7-Lunet-shNT or -shPI4KA cells stably expressing firefly luciferase were subcutaneously injected into nude mice and IVIS Spectrum *in vivo* imaging was performed at day 0, 3, 5, 10, 14, 18 and 21. 8 mice were used for each treatment group.

To further corroborate the importance of PI4KIIIα in liver pathogenesis *in vivo*, Huh7- Lunet-shNT and –shPI4KA cells stably expressing luciferase were transferred into SCID mice (Fig. S4G). Overall, larger areas of luciferase signal-positive cells were detected in the case of high PI4KIIIα expression (shNT) and some of the mice developed tumors (Fig. 4E). Signals of *PI4KA* knockdown cells, on the other hand, rapidly declined and most of them were below the threshold of detection on day 18 post transplantation (Fig. 4E). These data suggested that *in vivo* liver cancer cells with high PI4KIIIα expression had more capacity to form tumors than with low PI4KIIIα expression.

### Identification of PIK3C2γ as a downstream effector of PI4KIIIα

Next we aimed to understand whether the cytoskeleton reorganization was modulated directly by PI4P or by an alternative phosphatidylinositol phosphate upon conversion of PI4P, namely phosphatidylinositol 4,5-bisphosphate [PI(4,5)P2], phosphatidylinositol (3,4,5)-trisphosphate [PI(3,4,5)P3] and PI(3,4)P2. To this end, we employed shRNAs targeting all the members of the phosphatidylinositol-4-phosphate 5-kinases (PIP5Ks), class I phosphoinositide 3-kinases (PI3Ks) catalytic subunits, class II PI3Ks and PI4Ks protein families potentially contributing to the production and conversion of PI4P (Fig. S5A) [19]. Knockdown efficiencies ranged from 50% (shPIK3C2G) to 88% (shPIK3CD) (Fig. 5A). Silencing the other three PI4Ks members neither caused morphological changes, nor reduced phosphorylation levels of paxillin and cofilin, indicating that the regulation of this pathway was specific for PI4P originating from PI4KIIIα (Fig. 5B, 5C and S5B). Furthermore, neither silencing of PIP5Ks nor of class I PI3Ks catalytic subunits led to morphological changes and reduced p-paxillin and cofilin levels as observed for *PI4KA* knockdown (Fig. 5B and S5B), except for a moderate decrease in phosphorylation of the two proteins observed in *PIK3CD*-silenced cells (Fig. 5C). Interestingly, silencing of *PIK3C2G* showed comparable effects to *PI4KA* knockdown on both cell morphology and phosphorylation of paxillin and cofilin (Fig. 5B, 5C and S5B) and also recapitulated the effects of *PI4KA* knockdown on cell invasion and numbers of FAs (Fig. S5C and 5D-5F). Ectopic expression of *PIK3C2G* reverted the morphological changes in Huh7-Lunet- shPIK3C2G back to control status, indicating the specificity of the shRNA (Fig. S5D-E). Of note, depletion of *PIK3C2G*, paxillin, cofilin or other phospholipid kinases did not suppress HCV RNA replication as observed for *PI4KA* silencing (Fig. 5G-H and S5F-S5H). These data suggested that PIK3C2γ, encoded by *PIK3C2G* and converting PI4P to PI(3,4)P2, acted as a downstream kinase of PI4KIIIα to function on cytoskeleton-regulated cell morphology and invasiveness. In addition, the impact of PI4KIIIα activation on the cytoskeletal axis was likely uncoupled from its role in HCV RNA replication.

**Figure 5.**
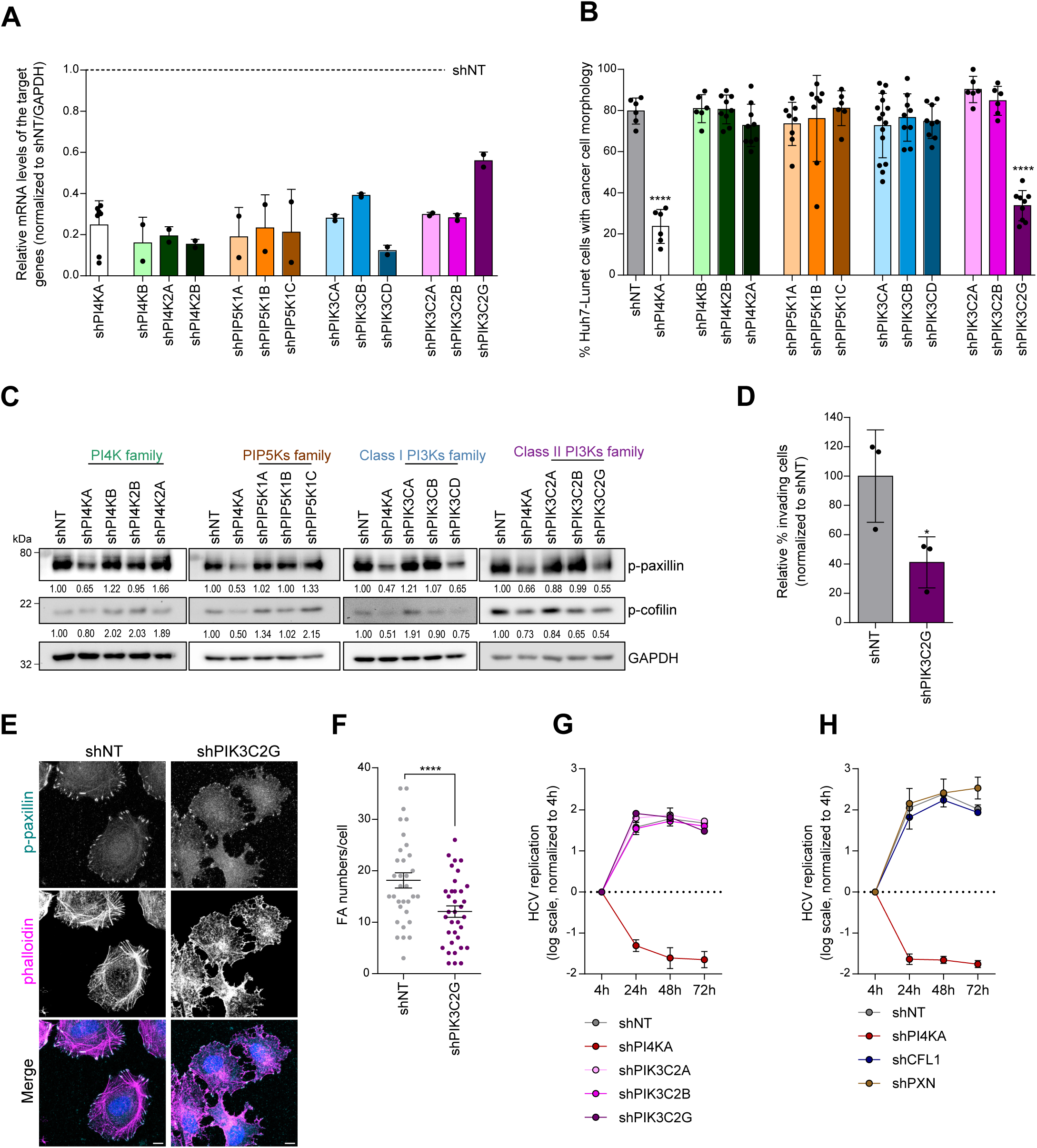
Identification of PIK3C2γ as a downstream partner of PI4KIIIα. (A-C) Silencing efficiency (A), cancer cell morphology (B) and levels of p-paxillin and p-cofilin (C) in Huh7-Lunet cells expressing the indicated shRNAs. Note that *PIK3CG* was not detected by RT-qPCR in Huh7-Lunet cells. Numbers below each lane represent relative expression levels compared to shNT. (D-F) Relative cell invasion rates (D), FAs staining (indicated by p-paxillin) (E) and numbers of FAs per cell (F) in Huh7-Lunet-shNT and shPIK3C2G. (G-H) Replication efficacy of HCV subgenomic replicons in Huh7-Lunet cells expressing the indicated shRNAs.

### PI(3,4)P2 pools at the PM are dependent on PI4KIIIα expression and activity

We next aimed to understand how PI(3,4)P2 distribution and abundance was regulated by PIK3C2γ and PI4KIIIα in liver cells. Huh7-Lunet cells showed diffuse cytoplasmic distribution of PI(3,4)P2 (Fig. 6A) and large clusters of PI(3,4)P2 at the sites of protrusions from the PM revealed by PM specific staining [20] (Fig. 6B), reminiscent of invadopodia [21] which are rich in actin and utilized by cancer cells to degrade the extracellular matrix for their invasiveness. Interestingly, PI(3,4)P2 enriched structures were rarely observed in *PIK3C2G* knockdown cells (Fig. 6B-C), whereas intracellular abundance of PI(3,4)P2 was slightly increased (Fig. 6A and 6C). These data suggested that in our HCC-derived cell culture model, PIK3C2γ contributed to the PI(3,4)P2 pools at the PM which were likely linked to invadopodia formation and cell invasion. In agreement with this finding, we found a concomitant reduction of PM-associated PI(3,4)P2 containing structures in *PI4KA* knockdown cells or cells treated with PI4KIIIα inhibitor (Fig. 6D-6F), indicating that a fraction of PI4P synthesized by PI4KIIIα was subsequently converted to PI(3,4)P2 at the PM by PIK3C2γ. Moreover, *PI4KA* silencing did not lead to any significant changes in the abundance of PI(4,5)P2 and PI(3,4,5)P3 at the PM (Fig. S6A-C), suggesting that PI4P synthesized by PI4KIIIα contributed exclusively to PI(3,4)P2 pools at the cell PM.

**Figure 6.**
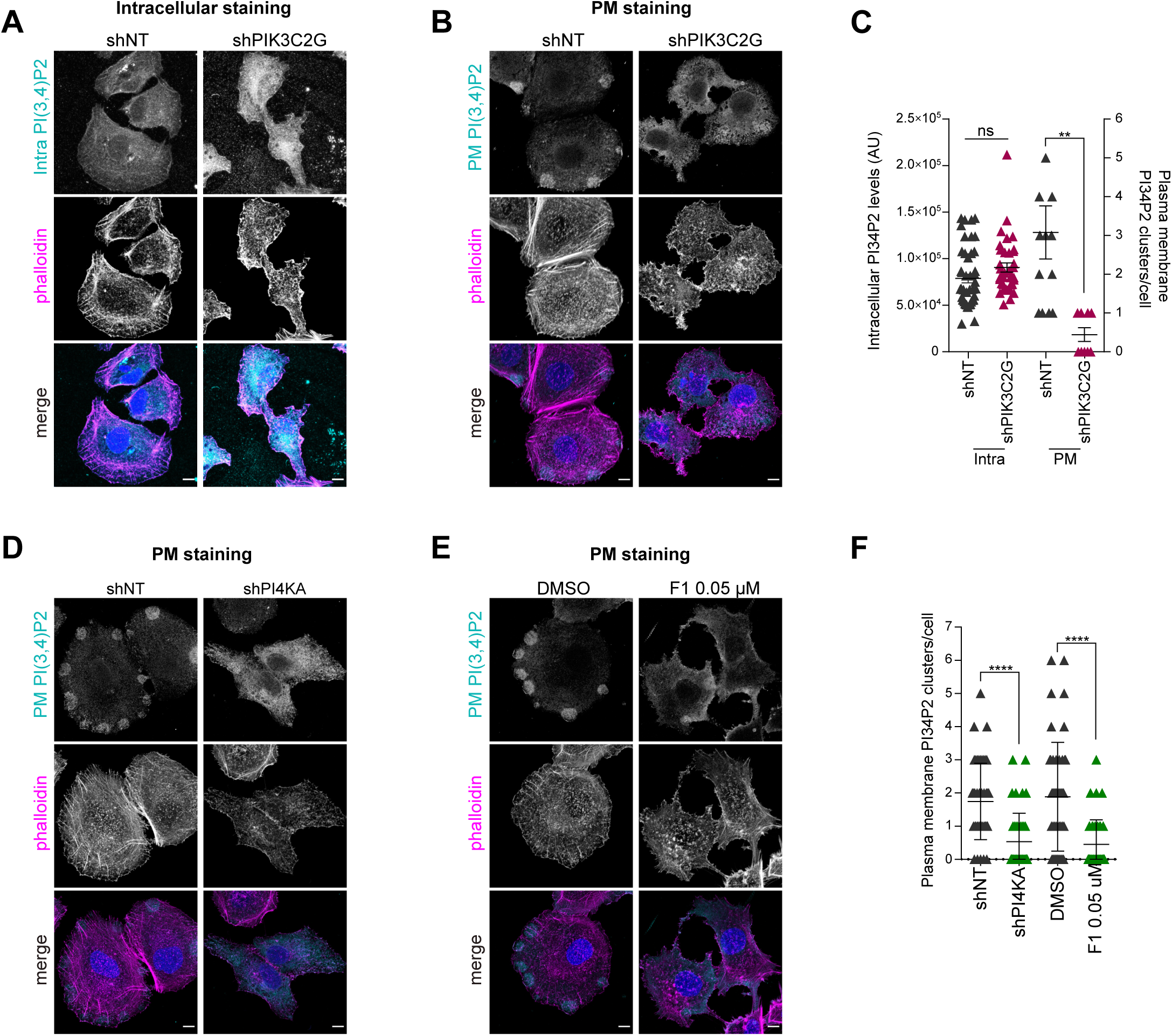
PI(3,4)P2 pools at the cell PM are dependent on PI4KIIIα expression and activity. (A-F) Huh7-Lunet cells expressing shPIK3C2G (A-C) or shPI4KA (D, F) or treated with PI4KA-F1 (E, F) were stained for phalloidin, DAPI and PI(3,4)P2, either intracellularly (A) or at the PM (B, D, E) and compared to shNT (A-D, F) or DMSO (E, F) treatment. Quantifications were performed in at least 10 cells (C, F).

### Akt2 is the mediator linking signaling from PI4KIIIα, PIK3C2γ to paxillin and further downstream

To identify kinase(s) downstream of PI(3,4)P2 involved in regulation of paxillin and cofilin phosphorylation, a phospho-kinase array was applied (Fig. 7A-B and S7A-B). Phosphorylation levels of Akt1/2/3 at Ser 473/474/472 were reduced 2-fold in cells depleted for *PI4KA* or *PIK3C2G* (Fig. 7A-B). In contrast, phosphorylation of focal adhesion kinase FAK and protein tyrosine kinase Src, often involved in paxillin phosphorylation [22], remained unchanged (Fig. 7A-C). For most other proteins, signal intensities were similar among the samples or were moderately altered in only one of the two knockdown samples (Fig. 7A-B, S7A-B). Western blot (WB) analysis confirmed that silencing of *PI4KA* and *PIK3C2G* reduced phosphorylation of both Akt1 and Akt2, but the effects on Akt2 were much stronger (Fig. 7C-D). While Akt1 was shown to be activated by HCV [23-25], there is so far no report of viral effects on Akt2. In fact, p-Akt2 S474 levels were strongly upregulated in HCV-infected samples (Fig. 7E). Furthermore, only HCV NS3-5B wt but not the mutants increased Akt2 phosphorylation (Fig. 7F), indicating that the Akt2 activation was associated with increased PI4KIIIα activity. Immunofluorescence using the PM specific staining protocol revealed that Akt2 was enriched at the periphery of the PI(3,4)P2 containing structures (Fig. 7G), supporting our hypothesis that Akt2 was the effector kinase recruited and activated by PI(3,4)P2. SC79, a pan-Akt activator, restored the p-Akt2 level (Fig. 7H), reverted the reduction of p-paxillin and p-cofilin (Fig. 7H) and in turn induced changes of cell morphology back to control status (Fig. 7I and S7C) in cells treated with PI4KA-F1. In addition, an Akt2 constitutively active mutant significantly reverted the morphological changes occurring in cells with low PI4KIIIα expression towards the cancer cell morphology (Fig. 7J and S7D) and restored their invasive properties (Fig. 7K). These data suggested that PI4KIIIα regulates cytoskeleton organization via PIK3C2γ/Akt2. Activation of Akt2 stimulates phosphorylation of paxillin and cofilin, controlling cell morphology and favoring cell motility of liver cancer cells.

**Figure 7.**
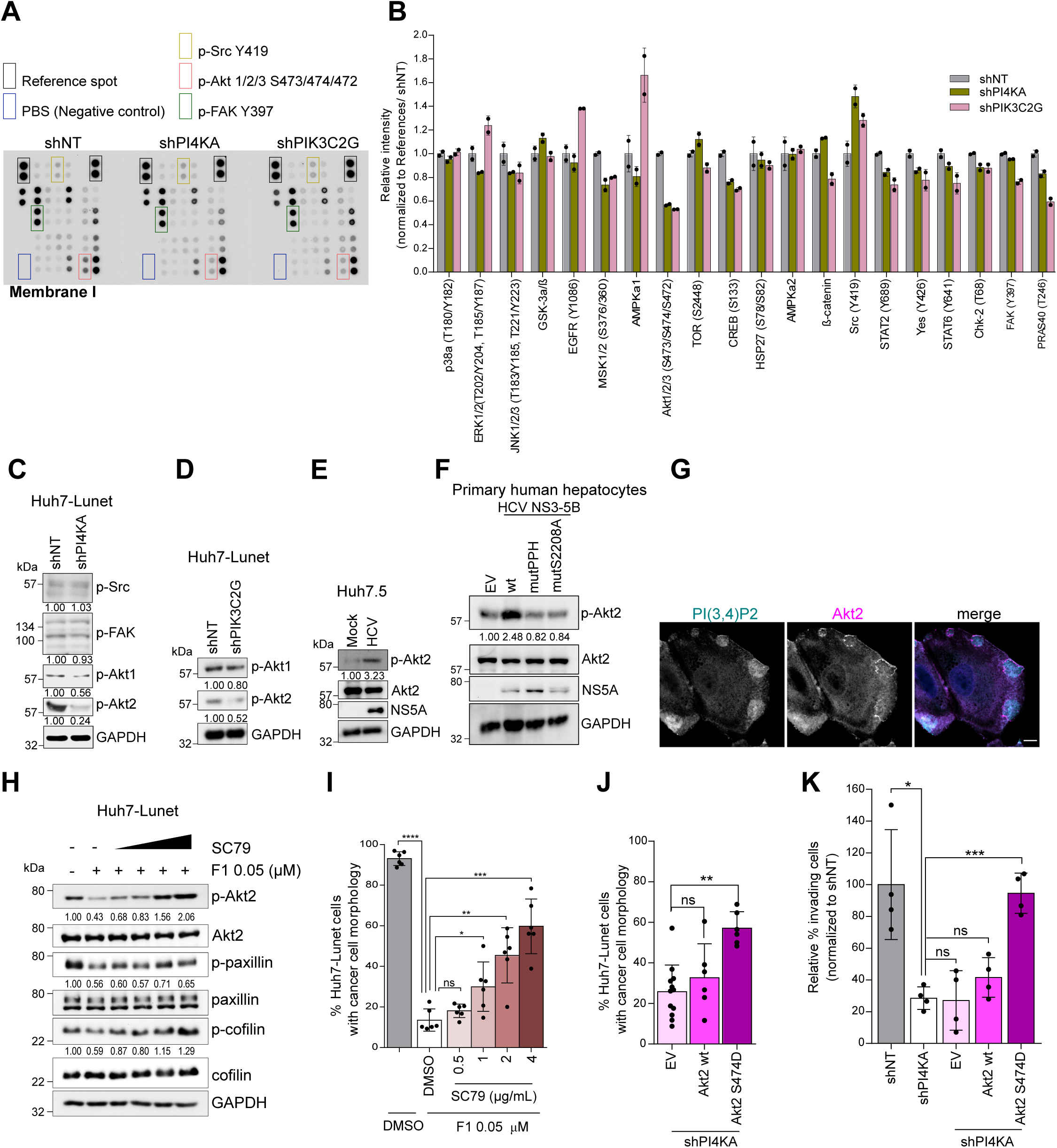
Akt2 is the mediator linking signaling from PI4KIIIα, PIK3C2γ to paxillin and further downstream. (A-B) Protein lysates from Huh7-Lunet expressing the indicated shRNA were incubated with the membrane I of the Proteome Profile human phospho kinase array kit (A). Relative signal intensities of each protein were quantified (B). (C-D) p-Akt1 and p-Akt2 levels in Huh7-Lunet-shNT versus -shPI4KA (C) or versus - shPIK3C2G (D). Akt3 was excluded due to its expression restricted to the brain and a few other organs . (E-F) p-Akt2 levels in HCV-infected Huh7.5 cells (E) and PHH expressing the indicated NS3-5B variants. (G) PM staining of PI(3,4)P2 and Akt2 in Huh7-Lunet cells. (H-I) Levels of p-Akt2, p-paxillin and p-cofilin (H) and cancer cell morphology (I) in Huh7- Lunet cells treated with PI4KA-F1 for 48 h, followed by SC79 for 30 min.. (J-K) Cancer cell morphology (J) and relative cell invasion rate (K) in Huh7-Lunet-shPI4KA expressing the indicated Akt2 variants. Numbers below each WB lane represent relative expression levels compared to the respective control shNT, mock or EV.

## DISCUSSION

PI4KIIIα is expressed at low abundance in hepatocytes [9, 10], but highly upregulated in HCC and associated with shorter disease-specific survival [9]. Here we show that high expression and/or activity of PI4KIIIα is important for migration and invasion of liver cancer cells via remodeling of the cytoskeletal network. Both *in vitro and in vivo* PI4KIIIα expression and activity were correlated with phosphorylation levels of paxillin and cofilin, essential factors for cancer cell motility [12-15], in agreement with previous studies associating paxillin and cofilin expression and/or activity with migration, invasion and metastasis of hepatocellular carcinoma [26, 27]. Still, expression levels and phosphorylation of additional cytoskeletal molecules and kinases were changed upon modulation of PI4KIIIα activity, suggesting that other cytoskeletal components cooperatively facilitate motility of HCC cells. Overall, our data might explain the poor prognosis of HCCs with high PI4KIIIα expression. However, animal models appropriately reflecting metastasis in liver cancer will be required to ultimately prove the contribution of PI4KIIIα to HCC progression.

Invasiveness and cytoskeletal changes of all HCC-derived cell lines included in our study were modulated by increased expression or activity of PI4KIIIα, pointing to the concentration of PI4P as a driving force mediating the phenotype. By targeting phospholipid kinases depending on PI4P as a substrate, we identified PIK3C2γ as the sole factor phenocopying PI4KIIIα, thus its phosphoinositide product PI(3,4)P2 appeared as a second messenger in this signal transduction pathway. PI(3,4)P2 so far has been considered as of minor importance, but recent studies have shown its relevance for cell invasion in several models by exerting important functions on cytoskeletal remodeling [28], membrane ruffle and podosome formation [29], and lamellipodia formation and maturation [30]. PIK3C2γ is mainly expressed in the liver with some traces found in the pancreas [31, 32]. Its expression is increased during liver regeneration after partial resection [32] and PIK3C2γ-deficient mice are viable and mainly have a defective insulin response, resulting in age dependent insulin resistance, obesity and fatty liver [31]. Potential roles of PIK3C2γ in cancer are particularly ill-defined. While some studies in murine pancreatic cancer models even regarded it as a tumor suppressor [33, 34], mutations in *PIK3C2G* were associated with poor prognosis in intrahepatic cholangiocarcinoma [35]. Our data now provide a first hint towards a contribution of PIK3C2γ to invasiveness of HCC. Since previous studies showed an increased expression of PI4KIIIα also in non-hepatic tumor tissues with migratory and invasive properties, promoting progression of renal [36], prostate [37] and pancreatic [38] cancer, increased PI4P concentration appear to drive similar processes via PIK3C2γ-independent mechanisms in other tissues.

The PI3K/Akt pathway is associated with multiple cancers, however, most studies are related to Akt1 and class I PI3Ks. Akt2 is the most abundant Akt isoform in the liver and its activity is correlated with the HCC incidence in c-Met-transfected mice [39]. While the specific binding of Akt2 to PI(3,4)P2 is well established [40] and its activation by PIK3C2γ in the context insulin signaling has been described previously [31], we identified a previously uncharacterized, liver specific link between PI4KIIIα/PIK3C2γ and Akt2 in cytoskeletal organization and cell invasiveness. While multiple classes of Akt inhibitors have already been established and evaluated preclinically and clinically, their impact in monotherapy is limited [41]. In general, Akt isoforms share redundant functions, but also have specific roles. Akt2 deficient mice are principally viable and develop diabetes [42], in line with the PIK3C2γ-dependent function in insulin signaling. However, specific targeting of Akt isoforms has been proven challenging [41]. In contrast, several highly potent and specific inhibitors of PI4KIIIα have been identified [8, 43], including the one used in our study. These molecules have not been further developed, mainly due to lethality of constitutive knockout and severe phenotypes of conditional knockout in mice [8, 43]. Therefore, PIK3C2γ could be a promising therapeutic target to prevent progression of liver cancer. In fact, first lead compounds with high specificity have been identified and tested *in vitro* on purified enzymes [44], but await further validation in cell culture models and *in vivo*.

Interestingly, not only increased PI4KIIIα expression, but also HCV infection is associated with poor prognosis and metastasis in HCC [9, 45]. It is therefore tempting to speculate, that higher aggressiveness of HCV-associated HCC is mediated by activation of PI4KIIIα, resulting in increased abundance of PI4P in HCV-infected cells [46]. Indeed, HCV infection or expression of NS3 to NS5B, critically involved in PI4KIIIα activation [46], promoted paxillin and cofilin phosphorylation and cell invasiveness in all cell culture models, including PHH, as well as *in vivo*, therefore phenocopying high PI4KIIIα abundance. However, PI4KIIIα activation by HCV is only crucial for HCV replication in human hepatocytes to generate sufficient amounts of PI4P, but rather deleterious in HCC derived cell lines with high PI4KIIIα abundance [10]. In line with these data, HCV replication is lower in HCC *in vivo* than in the surrounding tissue [47]. It seems therefore less likely that HCV directly promotes metastasis in HCC, but rather contributes to HCC severity by more indirect mechanisms, e.g. involving inflammatory responses [45, 48].

In conclusion, our study sheds light on a mechanistic connection of PI4KIIIα and liver pathogenesis. We suggest that PI4KIIIα upregulation in liver cancer cells or activation in response to HCV infection drives the progression of cancer by manipulating cytoskeletal components to promote migratory and invasive properties, via PIK3C2γ and Akt2. Pharmacological targeting of PI4KIIIα, PIK3C2γ or Akt2 may therefore serve as a therapeutic option to slow down liver cancer progression.

## CONFLICT OF INTEREST

The authors declare no competing interests.

## DATA AVAILABILITY

All data needed to evaluate the conclusions in the paper are present in the main text and/or the Supplementary information. Further data supporting the findings of this study isavailable upon request. Please contact the corresponding author.

## FINANCIAL SUPPORT STATEMENT

This project was funded by grants from the Deutsche Forschungsgemeinschaft Project Number 314905040-SFB/TRR209 to Vo.L., R.B., F.K., Pe.S., D.F.T., K.B., M.H.; Project Number 278191845 to Vo.L.; Project Number 272983813 to Vo.L. and R.B.. D.F.T. is supported by an ERC Starting Grant ‘CrispSCNAs’ (grant number: 948172) by the European Research Council. P.M. was supported by grants from the Ghent University Special Research Fund (UGent BOF) as well as an Excellence of Science grant (VirEOS2.0) from the Research Foundation – Flanders (FWO-Vlaanderen) and FNRS.

F.K. was supported by Deutsche Krebshilfe 70113873.

## AUTHOR CONTRIBUTIONS

Conceptualization: C.S.T. and Vo.L., Formal Analysis: C.S.T., T.P. and Vi.L., Funding acquisition: Vo.L., Investigation: C.S.T., J.K., M.B., J.Y., T. P., O.C., T.R., S.F., S.D., L.V., T.F.L, M.L., Ph.S., Methodology: C.S.T., T.P., F.K., K.B., P.M., M.H., M.S., Vi.L., D.F.T., Vo.L., Project administration: Vo.L., Resources: F.W.R.V., F.K., P.M., Pe.S., M.H., M.S., R.B., Vi.L., Software: C.S.T. and Vi.L., Supervision: Vo.L., Validation: C.S.T., J.K., Vo.L., Visualization: C.S.T., T.F.L., Vi.L., Writing – original draft: C.S.T. and Vo.L., Writing – review & editing: All authors.

## ACKNOWLEDGEMENTS

The authors would especially like to thank Ulrike Herian, Rahel Klein, Kai Volz, Lena Wendler, Jennifer Schmitt, Danijela Heide, Christine Schmitt, for excellent technical assistance. We thank Rani Burm, Bettina Fleischmann-Mundt and Peter Malin for their help with the mice experiments and Veronika Eckel for her assistance with scanning immunohistochemistry images. We are grateful to Takaji Wakita for the JFH-1 isolate, Janos Botyanszki for compound PI4KA-F1, Richard Harbottle for the pS/MARt vectors, Curt. C. Harris for HHT4 cells and Charles M. Rice for Huh7.5 cells and the 9E10 antibody. We also thank the Infectious Disease Imaging Platform (IDIP) headed by Vibor Laketa for facility use and help with microscopy.

## SUPPLEMENTAL MATERIALS AND METHODS

### PHH

PHH were obtained from the Department of General, Visceral, and Transplant Surgery at Hannover Medical School. PHH were processed from donors undergoing partial hepatectomy following written informed consent (approved by the ethic commission of Hannover Medical School/Ethik-Kommission der MHH, #252-2008). PHH isolation was performed using a 2-step collagenase perfusion technique as previously reported [1]. In brief, liver specimen were cannulated and flushed with pre-warmed (37 °C) washing buffer containing 2.5 mM EGTA. Thereafter, recirculating perfusion with pre-warmed (37 °C) digestion buffer containing 0.05% collagenase (Roche, Mannheim, Germany) was initiated. Upon sufficient digestion, the tissue was mechanically disrupted and the resulting cell suspension poured through a gauze-lined funnel followed by centrifugation (50 xg, 5 min, 4 °C). The cell pellet then was washed twice using ice-cold PBS (50 xg, 5 min, 4 °C) and resuspended in William’s medium E (all Biochrom AG, Berlin, Germany) supplemented with 1 µM insulin, 1 µM dexamethason/fortecortin, 100 U/mL penicillin, 100 µg/mL streptomycin, 1 mM sodium pyruvate, 15 mM HEPES buffer, 4 mM L-glutamine and 5% FCS. Cell number and viability were determined by the Trypan blue exclusion test. PHH were plated in 6-well culture dishes at 1.5×10^6^ cells/well.

### Plasmid constructs

Using Katahdin algorithms, a 97-mer oligonucleotide containing microRNA-based shRNA targeting sequence (see Table S2) was designed against each corresponding gene. In order to clone into pAPM vector, XhoI and EcoRI sites were introduced to flank sense and antisense sequence respectively as followed: for sense primers, the sequence tcgagaaggtatat was added in 5’ of the 97-mer oligonucleotides and the last a was deleted; while anti-sense primers was designed by reversing the sequence of the oligonucleotide and adding aat in 5’ and atataccttc in 3’. Each pair of sense and antisense primers was annealed by boiling at 95 °C for 5 min, followed by 30 min incubation at RT and was 5’ phosphorylated using T4 PNK (New England Biolabs) to create corresponding insert suitable for cloning into pAPM vector. Non targeting shRNA and shPI4KA were described previously [2] with the targeting sequences provided in the Table S2.

Construction of pWPI_PI4KA wt and pWPI_PI4KA D1957A with silent shRNA escape mutations were described elsewhere [3]. *PIK3C2G*, *PXN* (encodes paxillin) and *CFL1* (encodes cofilin) genes were amplified and introduced into AsiSI and AscI sites of the pWPI vector using the primers listed in the Table S3. The nucleotide exchange T91G was introduced to clone the constitutive active mutants *PXN* Y31D. In the case of *CFL1* S3D, the following nucleotide exchanges were introduced: T7G and C8A. All the mutations were introduced by overlap extension PCR. For the construction of pCSMB_ and pS/MARt_PI4KA wt, PI4KA D1957A, HCV NS3-5B wt, and HCV NS3-5B mutPPH, the respective genes of interest were amplified and introduced into NcoI and EcoRI sites of the sleeping beauty transposon vector pCSMB or NheI and BstZ17I sites of the pS/MARt vector using the primers listed in the Table S3.

### Transient HCV replication assay

Transient HCV replication was carried out as previously described [4] with minor modifications. In brief, replicon encoding plasmid DNA pFK-I341PI-Luc/NS3-3′/JFH1 with all HCV parts derived from HCV isolate JFH1 (genotype 2a) (in which its replication amplifies to high levels without the need for cell culture adaptation but is highly dependent on PI4KIIIα expression) and harboring hepatitis delta virus ribozymes was used for *in vitro* transcription. 2.5 µg of purified RNA was electroporated into 2×10^6^ Huh7-Lunet cells. Cells were then resuspended in 6 mL DMEM complete supplemented with 10 mM HEPES. 2 mL aliquots were seeded into each well of a 6-well plate for 4 h and 24 h time points, and 1 mL for 48 h and 72 h time points. Replication was determined by measuring firefly luciferase activity at the indicated time points post-electroporation and was normalized for transfection efficiency to the value obtained at 4 h time point.

To quantify luciferase reporter assay, cells were lysed in 350 µL Fluc lysis buffer (1% Triton X-100, 25 mM Glycine-Glycine (pH 7.8), 15 mM MgSO4, 4 mM EGTA (pH 7.8), 10% glycerol and 1 mM DTT). 100 µL of cell lysate was then mixed with 360 µL Fluc assay buffer (15 mM K3PO4 (pH 7.8), 25 mM Glycine-Glycine (pH 7.8), 15 mM MgSO4, 4 mM EGTA (pH 7.8), 1 mM DTT and 2 mM ATP). 200 µL luciferin (200 nM in 25 mM Glycine-Glycine) was then injected and measured for 20 sec in a tube luminometer (Lumat LB 9507, Berthold Technologies, Freiburg, Germany).

### Production of cell-culture-derived virus and infection of Huh7.5 with cell-culture-derived virus

HCVcc was produced according to the protocol published previously [5] with some modifications. In brief, 6×10^6^ Huh7.5 cells was resuspended in 400 µL of Cytomix buffer (120 mM KCl, 0.15 mM CaCl2, 10 mM K2HPO4/KH2PO4 (pH 7.6) and 25 mM HEPES) supplemented with 2 mM ATP and 5 mM glutathione. The cells were then mixed with 10 µg of *in vitro* transcribed HCV Jc1 viral RNA and electroporated using a Gene Pulser Xcell instrument (Bio-Rad Laboratories, Hercules, CA) in a 4-mm-gap cuvette. Cell culture supernatant collected at 4 days post electroporation was filtered through a 0.45 µm filtered and used as virus stock and stored at -80 °C. Virus titer was determined using method described by Lindenbach [6] which based on NS5A expression via immunohistochemistry.

One day prior to infection Huh7.5 cells were seeded on 10 cm dish. Cells was inoculated with HCVcc at MOI 0.5 for 6 h. Cells were then washed twice with PBS and then DMEM complete was replaced.

### Adenovirus mediated expression

HCV JFH NS3-5B expressions were achieved via adenovirus transduction which is generated according to the protocol from Agilent Technologies with some modifications. In brief, HCV NS3-5B wt and mutants (S232A and PPH) were cloned into pShuttle-CMV vector. The plasmids were then linearized with PmeI and then 1 µg of the purified DNA was electroporated into 50 µL BJ5183 electroporation competent cells bearing pAdEasy- 1 supercoiled vector to allow recombination occurs. Expected p-Ad vectors bearing HCV NS3-5B were then linearized with PacI. 5 µg of the corresponding linearized plasmid was transfected into 7×10^5^ 293T cells per 60-mm dish and incubated for 8 days. Cells were collected and resuspended in serum-free DMEM. Cell suspension was then subjected to four rounds of freeze/thaw by alternating the tubes between the -80 °C freezer and the 37 °C water bath with vortexing briefly after each thaw. Supernatant collected after centrifugation at 12000 xg for 10 min was used as primary viral stock and used to amplified virus titer by infection fresh HEK293T cells, followed by similar procedures for collecting virus. Target cells were transduced with virus stock and media was replaced after one day, followed by another 2 days of incubation.

### RT-qPCR

To quantify relative level of cellular mRNA, total RNA was extracted using a NucleoSpin RNA II kit (#740955, Macherey Nagel) according to instructions from the manufacturer. Complementary DNA (cDNA) was then synthesized using a High Capacity cDNA Reverse Transcription kit (#4368814, Applied Biosystems) and was diluted 1:10 with water before subjecting to RT-qPCR. The sequences of primers for cellular targets were obtained from the PrimerBank (Massachusetts General Hospital, The Center for Computational and Integrative Biology and Harvard Medical School) and were listed in the Table S3. RT-qPCRs were performed in triplicate using an iTaq Universal SYBR Green kit (#1725121, Bio-Rad) with a Bio-Rad CFX96 Touch Real-Time PCR system, and data were analyzed using the corresponding software.

### Immunoblotting

Cells seeded in 6-well plates were washed twice with ice-cold PBS, harvested by lysis with 90 µL per well of lysis buffer containing 20 mM Tris HCl (pH 7.2), 150 mM NaCl, 10% glycerol, 1% NP-40, 10 mM NaF, 30 mM sodium pyrophosphate, 1 mM EDTA, 1 mM Na3VO4, 1 mM phenylmethylsulfonyl fluoride, and a protease inhibitor cocktail and incubated on ice for 20 min. Supernatant collected after centrifugation at 13500 rpm at 4°C was mixed with Laemmli 6X buffer (375 mM Tris.HCl, 9% SDS, 50% glycerol, 0.03% bromophenol blue and 9% β-mercaptoethanol) and was heated 5 min at 95°C and loaded onto polyacrylamide-SDS gel. After electrophoresis proteins were transferred to a PVDF membrane and was blocked in 5% milk (in PBS supplemented with 0.25 % Tween (PBS- T)) for 1 h. After over-night incubation of the membrane at 4°C with primary antibodies diluted in PBS-T supplemented with 1% bovine serum albumin (BSA), the membrane was washed 3 times for 15 min with PBS-T and incubated with secondary antibodies conjugated to horseradish peroxidase diluted in PBS-T supplemented with 1% milk for 1 h at RT. After extensive washing, bound antibodies were detected with the ECL Plus Western Blotting Detection System. Akt activator SC79 was obtained from Abcam (#ab146428) to evaluate the importance of Akt in PI4KIIIα-regulated cell morphology and phosphorylation of paxillin and cofilin.

### Immunofluorescence analysis and phosphoinositides quantification

Cells were firstly subjected to either intracellular or PM staining protocol.

#### Intracellular staining

Intracellular phosphoinositides, HCV NS5A, p-paxillin, α-tubulin and cytokeratin-8 were stained as described elsewhere [2]. In brief, cells seeded on coverslips were washed twice with PBS before fixation in 4% paraformaldehyde for 15 min. Cells were then permeabilized with 50 µg/mL digitonin (for staining phosphoinositides) or 0.1% Triton X- 100 for 10 min and blocked in 3% BSA for 30 min. Cells were incubated with primary antibodies diluted in 3% BSA either at RT for 2 h or at 4 °C overnight. Cells were further incubated with Alexa 488- or 568-conjugated secondary antibodies and/or rhodamine phalloidin in 3% BSA for 45 min at RT with a dilution of 1:500. Nuclei were stained using 4‘,6-diamidino-2- Phenylindole dihydrochloride (DAPI) for 1 min at a dilution of 1:4000. Cells were mounted with Fluoromount G.

#### PM staining

PM phosphoinositides were stained as in previous study [7]. In brief, after aspiring media, buffer A (4% paraformaldehyde and 0.2% glutaraldehyde in PBS) was added directly to allow fixation at RT for 15 min. After 3 times rinsing with buffer B (PBS containing 50 mM NH4Cl), slides with cover slips were place on ice. The cells were then blocked for 45 min with buffer C (5% (v/v) NGS, 0.5% saponin in buffer B), followed by primary antibodies in buffer D (5% NGS (v/v) and 0.1% saponin in buffer B) for 2 h. After 3 washes with buffer B, secondary antibodies prepared in buffer D were added and incubated for 45 min. Nuclei was stained with DAPI for 1 min, followed by 4 times rinsing with buffer C. Cells were then post-fixed with buffer E (2% paraformaldehyde in PBS) on ice for 10 min and at RT for 10 min. After 2 times rinsing with buffer B and one time with distilled water, cells were mounted with Fluoromount G.

Whole-cell z-stacks were acquired using a 63x oil immersion objective (NA, 1.4; Leica APO CS2) on a Leica TCS SP8 (Leica Microsystems) confocal laser scanning microscope. Quantification of intracellular and PM phosphoinositides signals was described before [2].

### Antibodies and molecular probes

Anti-PI4KIIIα (#4902, Cell Signaling), anti-GAPDH (#sc-47724, Santa Cruz), anti-paxillin (#05-417, Millipore), anti-paxillin phospho Y31 (#44-720G, Thermo Fisher Scientific), anti-cofilin (#5175, Cell Signaling), anti-cofilin phospho S3 (#3313, Cell Signaling), anti-Akt2 (#3063, Cell Signaling), anti-Akt2 phospho S474 (#8599, Cell Signaling), anti-Akt1 (#2938, Cell Signaling), anti-Akt1 phospho S473 (#4060, Cell Signaling), anti-E-cadherin (#3195, Cell Signaling), anti-α-tubulin (#A01410, GenScript), anti-NS5A (clone 9E10, gift from C.M. Rice, Rockefeller University, N.Y), anti-PI4P (#Z-P004, Echelon), anti-PI(3,4)P2 (#Z-P034, Echelon), anti-PI(4,5)P2 (#Z-P045, Echelon), anti-PI(3,4,5)P3 (#Z-P345,

Echelon), anti-Src phospho Y419 (#AF2685, R&D Systems), anti-FAK phospho Y397 (#3283, Cell Signaling), horseradish peroxidase conjugated anti-mouse IgG (#A4416, Sigma-Aldrich), horseradish peroxidase conjugated anti-rabbit IgG (#A6154, Sigma-Aldrich), Alexa Fluor 488 conjugated anti-mouse IgM (#A21042, Invitrogen), Alexa Fluor 488 conjugated anti-mouse IgG (#A21202, Invitrogen), Alexa Fluor 488 conjugated anti-rabbit IgG (#A-11008, Invitrogen), Alexa Fluor 568 conjugated anti-rabbit IgG2α (#A- 21134, Invitrogen), DAPI (#D21490, Invitrogen), rhodamine phalloidin (#R415, Invitrogen).

### Bright field microscopy

To obtain bright field microscopic images, cells were seeded in 6-well plate and cell morphology was monitored over time. Images were acquired 2-3 days post seeding when cells reach 50-75% confluence using an inverted phase contrast microscope (Leica DMIL LED Fluo, S/N 479642, Germany with a camera Leica MC170 HD #11541550) with 20x objective lens (N PLAN 20x/0.35 PH1, #11541550). Photos of at least six random areas covering minimum 200 cells in each condition were subjected for quantification.

### Machine learning- based automated cell morphology classification and analysis of histology sections

In order to quantify the prevalence of characteristic cellular phenotypes (cancer cell morphology versus compact morphology) across different experiments in a consistent and unbiased fashion we developed an AI-based classification workflow. The segmentation of the entire cells from the bright field microscopy images was not possible as the border between cells is not clearly distinguishable. Therefore, we decided to segment the cellular nuclei which are visually apparent in bright field images and there is typically one per cell. The nuclear area together with the neighboring pixels has sufficiently distinguishable features to be able to classify cells in cancer cell morphology and compact morphology phenotypes in the final classification step without considering the entire cellular area. To segment nuclei in the bright field images without the specific nuclear marker we employed an online deep learning platform Apeer from the company Zeiss (https://www.apeer.com) where we trained a model using the subset of our data and used it to segment the nuclei in all our bright field imaging data. The segmented nuclei channel together with raw bright field images were subsequently used in the software ilastik [8] to train a Random Forest classifier that is able to distinguish between cancer cell morphology and compact morphology phenotypes. The training set of data was arbitrary selected and very sparsely labelled (<1% of total objects were manually categorized into cancer cell morphology and compact morphology categories, object neighborhood (30px) was considered for classification). The same ilastik model was used for cell classification across all experiments, ensuring consistent application of criteria for cancer cell morphology and compact morphology categorization throughout the project.

To quantify the signal area on histology sections, we leveraged the pixel classification workflow of the ilastik software. We trained a model using arbitrary selected area of one histology section where we very sparsely manually assigned pixels (<0.1% of total pixels) into “signal” and “background” categories. The same model was used to quantify the signal area in histology sections in Figure 5C.

### Histopathology and evaluation

After overnight fixation in 10% buffered formalin, representative specimens of the liver were routinely dehydrated, embedded in paraffin, and cut into 4 µm-thick sections. Tissue sections were routinely stained with hematoxylin and eosin and Picrosirius red, respectively. All staining methods were conducted according to standard protocols.

Histopathological evaluation was performed using a semiquantitative scoring system [9] and adapted for the present mouse model. Comparative analysis includes steatosis, hepatocellular hypertrophy, fibrosis, inflammation, liver cell injury and bile duct proliferation. Scoring based on the evaluation of the whole tissue sections to compensate unavoidable variabilities of the cutting plane and the inhomogeneous distribution of the lesions. All tissue sections were analyzed blinded concerning genotype and treatments of the mice.

### Cell migration assay

The ibidi Culture-Insert 2 Well was used to evaluate cell migration to provide a defined size of the cell-free gap. During trypsinization, the inserts were plated into a 24-well plate. To minimize the effect of cell proliferation on the assay, cells were seeded at the density of 1.5×10^4^ to have a 100% cell confluency. At 4 h post seeding, images of the cell-free gaps were taken using phase contrast microscopy and served as control starting point of the assay. For each gap, images were taken at the two ends using an inverted phase contrast microscope (Leica DMIL LED Fluo, S/N 479642, Germany with a camera Leica MC170 HD, #11541550) with 4x objective lens (HI PLAN 4x/0.10, #506226) and the exact same gap position was monitored during the entire experiment. Cells were further monitored and images were then taken at 24 and 48h post seeding.

### Cell proliferation assay

Cell proliferation was determined via measuring cell viability at different time points after seeding (4 h, 24 h, 48 h, 72 h) using CellTiter-Glo Luminescent Cell Viability Assay (#G7571, Promega). In brief, 25 µL of CellTiter-Glo Reagent was added to 25 µL of medium containing cells for a 96-well plate, followed by 2 min mixing on an orbital shaker to induce cell lysis. After 10 min incubation at RT to stabilize luminescent signal, luminescence was recorded.

### Human phospho-kinase array

A human phosphokinase array (#ARY003B, Proteome Profiler, R&D systems) was applied using lysates from Huh7-Lunet cells stably expressing shNT, shPI4KA or shPIK3C2G according to the manufacturer’s instruction with few modifications. In brief, the cells cultured in 15 cm dishes were lysed in 500 µL lysis buffer 6, followed by 30 min rocking on ice. Supernatant was collected by a centrifugation at 14000 xg for 5 min and protein concentration was determined by Bradford assay. From each sample, 600 µg of total protein diluted in Array Buffer 1 were divided evenly into two parts and each was incubated with either membrane I or II (already blocked with Array Buffer 1) separately at 4 °C, overnight. After three washes with 1x Wash buffer, the membranes were incubated with the respective Detection Antibody Cocktail A or B for 2 h at RT. This was followed by another incubation with diluted Streptavidin-HRP (in 1x Array Buffer 2/3) and prepared Chemi Reagent Mix with three washes in between. The membranes were then treated with an ECL detection kit (PerkinElmer) and chemiluminescence was recorded using ECL imaging system (Intas).

### Cytoskeleton antibody array

To prepare for Cytoskeleton Phospho Antibody Array (#PCP141, Full Moon Biosystems), Huh7-Lunet cells were treated with 0.05 µM PI4KA-F1 or DMSO for 3 days. After 5 washes with cold PBS, cell pellets were collected by a centrifugation at 700 rpm, 5 min. Cell pellets were then transferred to Full Moon BioSystems for further assay. In brief, non-denaturing lysis buffer was used for protein extraction, followed by a biotinylation. These labeled samples were then incubated with antibody array and detection was facilitated by dye conjugated streptavidin.

### Mice experiments

#### Hydrodynamic tail-vein injection of plasmids expressing PI4KIIIα or HCV *nonstructural proteins*

The group size for individual animal experiments was determined on the basis of our experience with previous similar experiments. Hydrodynamic tail vein injection was performed in 7-8-week-old female C57Bl/6 mice (Janvier). For hydrodynamic tail vein injection and Sleeping Beauty transposase-mediated stable expression of constructs, a sterile 0.9% NaCl solution/plasmid mix was prepared containing 20 µg of respective plasmid (pCSMB-EV or –HCVNS3-5Bwt/mut or PI4KA wt/mut) together with 4 µg CMV- SB13 Transposase (1:5 ratio) per animal. For hydrodynamic tail vein injection and scaffold/matrix attachment region (pS/MARt)-based stable expression of constructs, a sterile 0.9% NaCl solution/plasmid mix was prepared containing 40 µg of respective plasmid (pS/MARt-EV or –HCVNS3-5Bwt/mut or PI4KA wt/mut) per animal. Mice were randomly assigned to experimental groups and injected with the 0.9% NaCl solution/plasmid mix into the lateral tail vein with a total volume corresponding to 10 % of body weight in 5-7 seconds. 4 weeks post-injection, mice were euthanized in compliance with all relevant ethical regulations determined in the animal permit. Livers were extracted and tissue samples were either fixed overnight in 4% PFA or snap-frozen at -80 °C for further use. All animal experiments were approved by the regional board Karlsruhe, Germany. The fixed samples were delivered to Center for Model System and Comparative Pathology, Heidelberg University Hospital for immunohistochemistry staining and images were scanned in the NCT-Gewebebank. Pathological status of the samples was further evaluated using HSE staining by T.P. For each frozen liver, several small pieces were taken and pooled together for a representative sample. Protein lysates were extracted using 200 µL lysis buffer with the aid of Dounce tissue grinder (Sigma Aldrich, D8938). Expression of p-paxillin, p-cofilin and GAPDH was assessed by western blot.

#### Injection of human liver cancer cells with modulated PI4KA expression into SCID mice

Huh7-Lunet cells were firstly transduced with adenoviruses expressing a firefly luciferase to favor *in vivo* monitoring of the injected cells and tumor formation. These cells were then either stably knocked down of *PI4KA* or control silenced using lentivirus-mediated gene transduction. Morphology of these cells as well as expression levels of PI4KIIIα was assessed to ensure an efficient silencing. All mice-related experimental procedures were conducted in accordance with the Federation of Laboratory Animal Science Associations (FELASA) and experiment procedures were granted by regional animal ethics committees (approvals Dnr 13278-2019). Four-week-old female athymic nude mice (Balb/c nu/nu) were obtained from Charles River Laboratories, Sulzfeld, Germany, and maintained under standard conditions at the preclinical laboratory (PKL), Karolinska University Hospital Huddinge, Sweden. All mice had free access to sterilized food and autoclaved water. Tumor inoculation was started after 1 week of acclimatization. On day 0, a suspension of cells (1 or 5×10^6^ cells in 0.2 mL phosphate-buffered saline) was injected subcutaneously into the dorsal flank. *In vivo* bioluminescent assays were performed on days 3, 5, 10, 14, 18 and 21. A dose of 150 mg luciferin/kg body weight of d-luciferin was injected intra- peritoneally (i.p.) 10-15 minutes before *in vivo* imaging. On day 21 the mice were sacrificed, and organs were collected including livers, lungs, spleens and tumors.

#### HCV infection of human-liver chimeric mice

Human liver-chimeric mice were generated by transplantation of primary human hepatocytes (donor L191501, Lonza, Switzerland) into hHomozygous uPA+/+-SCID mice as described before [10]. Mice with high-level humanization of the liver (human albumin level 2.5 to 5.6 mg/mL in mouse plasma) were transplanted with primary human hepatocytes. Infection of human-liver chimeric mice was achieved with HCV WT isolates from sera of infected patients gt1b, strain GLT1 [11] or gt1a, strain mH77c [12]. The mice were sacrificed at 8 weeks post infection. Expressions of human albumin, HCV NS5A and p-paxillin in mice livers were assessed by immunohistochemistry on consecutive slices. Briefly, 2 µm consecutive sections of mice liver were stained with antibodies against human albumin, HCV NS5A or p-paxilin using the BOND-MAX Automated IHC/ISH Stainer (Leica). All reagents used were purchased from Leica. Detection was performed using a secondary antibody-polymer (Leica) coupled to horseradish peroxidase.

### Cell lines

The following cell lines were used in this study: HEK293T MCB [13], Huh7 [14], Huh7- Lunet [15], Huh7.5 [16], HepG2 [17], Hep3B [18], HHT4 (Gift from Curt. C. Harris, Center for Cancer Research; National Cancer Institute, Maryland), Hep55.1C [19], Huh7-Lunet- shNT, -shPI4KA, -shPI4KB, -shPI4K2A, -shPI4K2B, -shPIP5K1A, -shPIP5K1B, - shPIP5K1C, -shPIK3CA, -shPIK3CB, -shPIK3CD, -shPIK3C2A, -shPIK3C2B, and - shPIK3C2G were created in this study.

## SUPPLEMENTAL TALBES

**Table S1.** Fold change of cytoskeleton proteins (phosphorylated and total forms) revealed by a Cytoskeleton Phospho Antibody Array (Full Moon Biosystems).

| Antibody List (phosphorylated protein) | Fold Change PI4KA-F1/DMSO | Antibody List (phosphorylated protein) | Fold Change PI4KA-F1/DMSO |
| --- | --- | --- | --- |
| Actin Pan (a/b/g) (Phospho-Tyr55/53) | 1.41 | MEK1 (Phospho-Thr286) | 1.17 |
| Calmodulin (Phospho-Thr79/Ser81) | 0.74 | MEK1 (Phospho-Thr291) | 1.08 |
| CaMK1-alpha (Phospho-Thr177) | 0.96 | MEK1 (Phospho-Ser298) | 0.47 |
| CaMK2 (Phospho-Thr286) | 1.18 | MEK2 (Phospho-Thr394) | 0.82 |
| CaMK2-beta/gamma/delta (Phospho-Thr287) | 0.59 | Merlin (Phospho-Ser10) | 1.31 |
| CaMK4 (Phospho-Thr196/200) | 1.01 | Merlin (Phospho-Ser518) | 1.14 |
| Cofilin (Phospho-Ser3) | 2.13 | MKK3/MAP2K3 (Phospho-Ser189) | 1.08 |
| Cortactin (Phospho-Tyr421) | 0.98 | MKK3/MAP2K3 (Phospho-Thr222) | 0.7 |
| Cortactin (Phospho-Tyr466) | 0.96 | MKK6 (Phospho-Ser207) | 0.96 |
| c-Raf (Phospho-Ser296) | 1.06 | MKK7/MAP2K7 (Phospho-Ser271) | 0.73 |
| c-Raf (Phospho-Ser43) | 1.11 | Myosin regulatory light chain 2 (Phospho-Ser18) | 0.92 |
| CrkII (Phospho-Tyr221) | 0.97 | p130Cas (Phospho-Tyr165) | 1.01 |
| ERK1-p44/42 MAP Kinase (Phospho-Thr202) | 0.98 | p130Cas (Phospho-Tyr410) | 0.99 |
| ERK1-p44/42 MAP Kinase (Phospho-Tyr204) | 1.03 | Paxillin (Phospho-Tyr118) | 1.02 |
| ERK3 (Phospho-Ser189) | 0.94 | Paxillin (Phospho-Tyr31) | 0.7 |
| Ezrin (Phospho-Tyr353) | 1.05 | PI3-kinase p85-subunit alpha/gamma (Phospho-Tyr467/Tyr199) | 0.66 |
| Ezrin (Phospho-Tyr478) | 0.99 | PKA CAT (Phospho-Thr197) | 1.31 |
| Ezrin (Phospho-Thr566) | 1.01 | PKC alpha (Phospho-Tyr657) | 1.12 |
| FAK (Phospho-Tyr397) | 0.92 | PKC alpha/beta II (Phospho-Thr638) | 1.17 |
| FAK (Phospho-Tyr407) | 1.36 | PKC pan activation site (Phospho) | 1.15 |
| FAK (Phospho-Tyr576) | 1.29 | PLC beta-3 (Phospho-Ser1105) | 1.25 |
| FAK (Phospho-Tyr861) | 1 | PLC beta-3 (Phospho-Ser537) | 0.96 |
| FAK (Phospho-Ser910) | 0.91 | Rac1/cdc42 (Phospho-Ser71) | 0.86 |
| FAK (Phospho-Tyr925) | 0.9 | Rho/Rac guanine nucleotide exchange factor 2 (Phospho-Ser885) | 0.77 |
| Filamin A (Phospho-Ser2152) | 1.24 | Src (Phospho-Tyr418) | 0.52 |
| Gab2 (Phospho-Ser159) | 0.97 | Src (Phospho-Tyr529) | 1.01 |
| GTPase activating protein (Phospho-Ser387) | 1.3 | Src (Phospho-Ser75) | 0.98 |
| LIMK1 (Phospho-Thr508) | 1.19 | VASP (Phospho-Ser157) | 1.02 |
| MEK1 (Phospho-Ser217) | 1.04 | VASP (Phospho-Ser238) | 1.02 |
| MEK1 (Phospho-Ser221) | 0.99 |  |  |

**Table S1 (continued)**
| **Antibody List (total protein)** | **Fold Change**<br>**PI4KA-F1/DMSO** | **Antibody List (total protein)** | **Fold Change**<br>**PI4KA-F1/DMSO** |
| --- | --- | --- | --- |
| Actin alpha-1 skeletal muscle (N-term) | 1.02 | MEK1 (Ab-217) | 0.82 |
| Actin alpha-2/3 (N-term) | 0.92 | MEK1 (Ab-221) | 0.96 |
| Actin Pan (a/b/g) (Ab-55/53) | 0.65 | MEK1 (Ab-286) | 0.69 |
| ACTN1 (F-Actin) (inter) | 0.81 | MEK1 (Ab-291) | 0.76 |
| Beta actin | 0.93 | MEK1 (Ab-298) | 0.65 |
| Calmodulin (Ab-79/81) | 1.3 | MEK2 (Ab-394) | 1.18 |
| CaMK1-alpha (Ab-177) | 1 | MEKKK 1 (inter) | 0.86 |
| CaMK1-beta (inter) | 0.81 | MEKKK 4 (inter) | 0.92 |
| CaMK2 (Ab-286) | 0.73 | Merlin (Ab-10) | 0.74 |
| CaMK2-beta/gamma (inter) | 0.88 | Merlin (Ab-518) | 0.79 |
| CaMK2-beta/gamma/delta (Ab-287) | 1.34 | MKK3/MAP2K3 (Ab-189) | 0.89 |
| CaMK4 (Ab-196/200) | 0.93 | MKK3/MAP2K3 (Ab-222) | 1.19 |
| CaMK5 (inter) | 0.77 | MKK6 (Ab-207) | 1.04 |
| Cofilin (Ab-3) | 0.5 | MKK7/MAP2K7 (Ab-271) | 1.11 |
| Cortactin (Ab-421) | 0.86 | Myosin regulatory light chain 2 (Ab-18) | 1.1 |
| Cortactin (Ab-466) | 0.79 | NCK2 (C-term) | 0.99 |
| c-Raf (Ab-296) | 0.92 | p130Cas (Ab-165) | 1.01 |
| c-Raf (Ab-43) | 0.82 | p130Cas (Ab-410) | 1.14 |
| CrkII (Ab-221) | 0.88 | Paxillin (Ab-118) | 0.92 |
| ERK1/2 (N-term) | 0.83 | Paxillin (Ab-31) | 1.17 |
| ERK1-p44/42 MAP Kinase (Ab-202) | 0.82 | PI3-kinase p85-subunit alpha/gamma (Ab-467/199) | 1.26 |
| ERK1-p44/42 MAP Kinase (Ab-204) | 0.82 | PIP5K (inter) | 0.83 |
| ERK3 (Ab-189) | 0.96 | PKA CAT (Ab-197) | 0.8 |
| Ezrin (Ab-353) | 0.96 | PKC alpha (Ab-657) | 0.87 |
| Ezrin (Ab-478) | 1.05 | PKC alpha/beta II (Ab-638) | 0.8 |
| Ezrin (Ab-566) | 0.86 | PKC pan activation site | 0.8 |
| FAK (Ab-397) | 0.89 | PLC beta-3 (Ab-1105) | 0.98 |
| FAK (Ab-407) | 0.83 | PLC beta-3 (Ab-537) | 0.93 |
| FAK (Ab-576) | 0.92 | Rac1/cdc42 (Ab-71) | 0.82 |
| FAK (Ab-861) | 1.06 | Rho/Rac guanine nucleotide exchange factor 2 (Ab-885) | 1.18 |
| FAK (Ab-910) | 0.77 | Src (Ab-418) | 1.04 |
| FAK (Ab-925) | 0.9 | Src (Ab-529) | 0.9 |
| Filamin A (Ab-2152) | 0.72 | Src (Ab-75) | 1.16 |
| Gab2 (Ab-159) | 1.04 | VASP (Ab-157) | 0.76 |
| GAPDH | 0.75 | VASP (Ab-238) | 0.88 |
| GTPase activating protein (Ab-387) | 0.65 | WASP (Ab-290) | 0.99 |
| LIMK1 (Ab-508) | 0.9 | WAVE1 (Ab-125) | 0.98 |
| LIMK1/2 (Ab-508/505) | 0.85 |  |  |

**Table S2.**
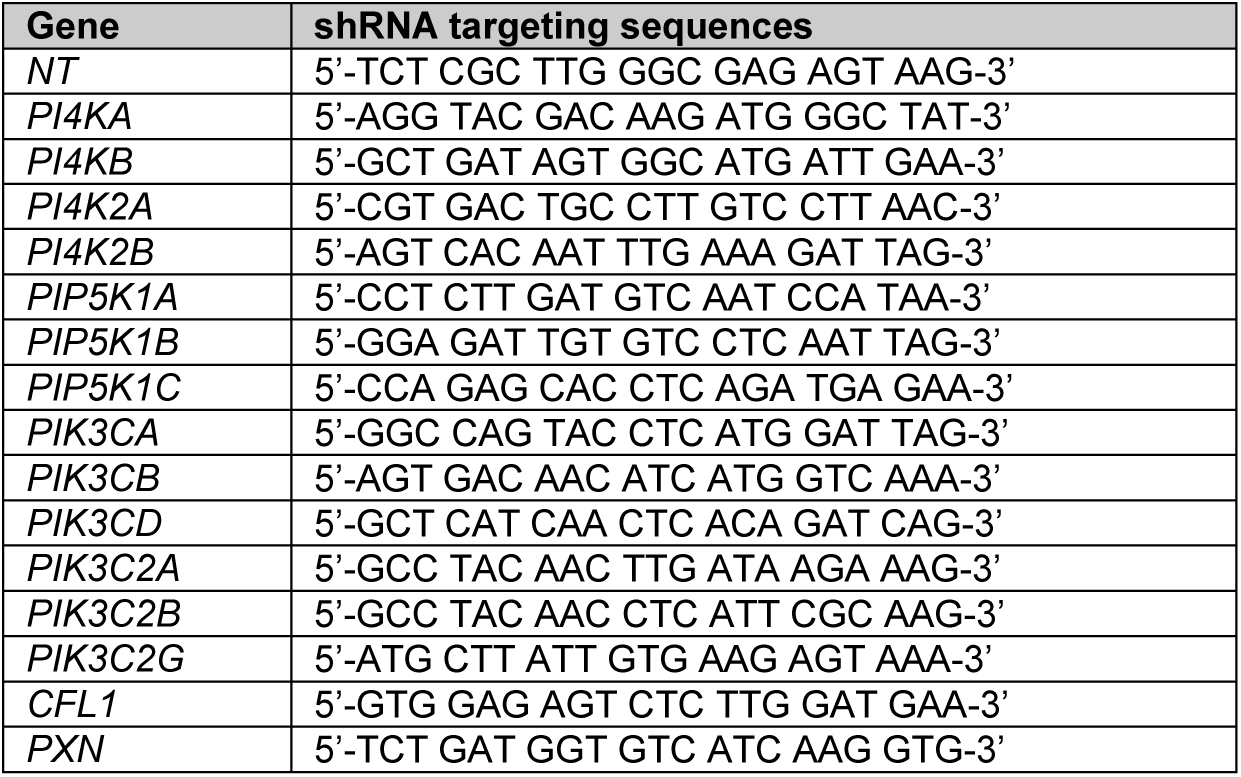
shRNA targeting sequences used in this study.

**Table S3.**
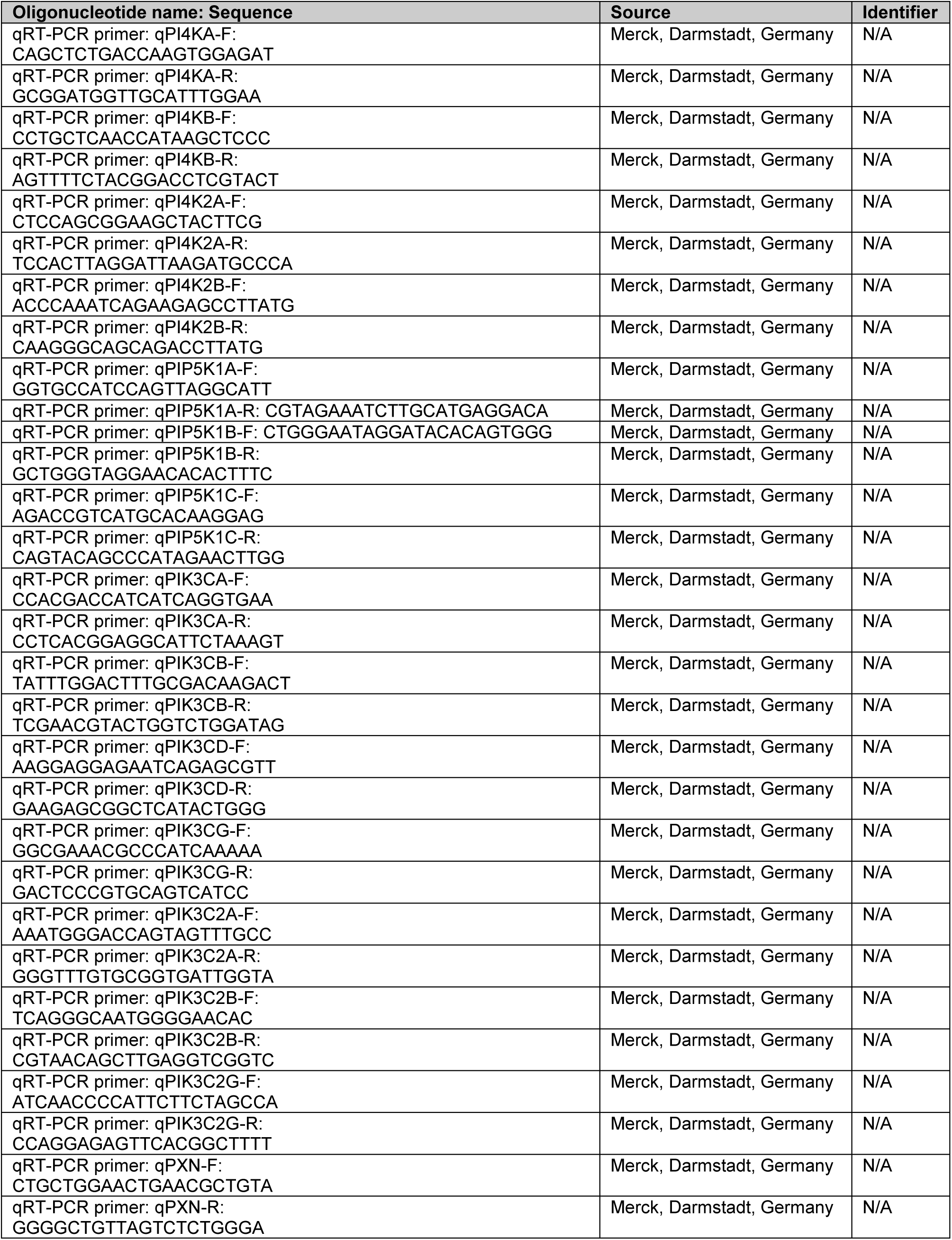
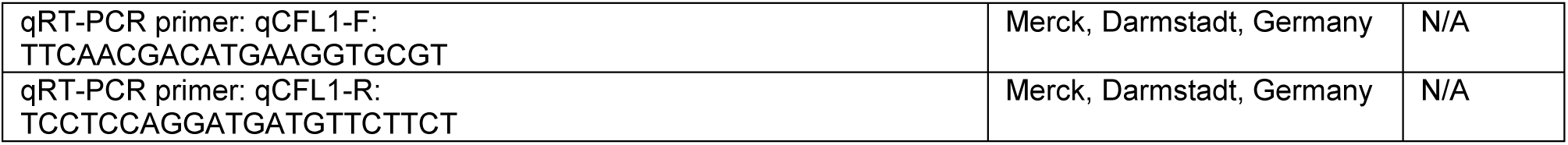
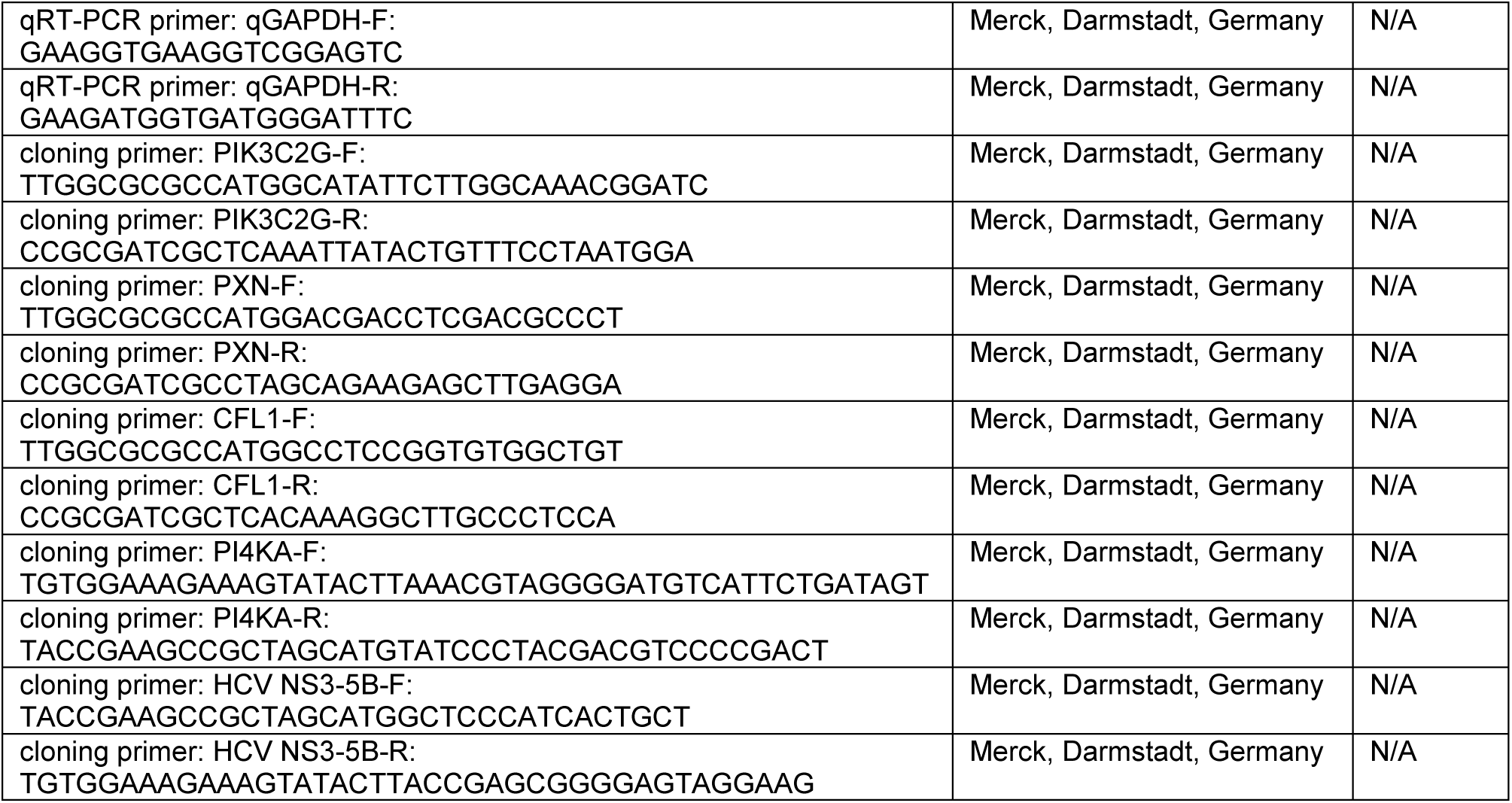
Oligonucleotides sequences used in this study.

## SUPPLEMENTAL FIGURES

**Figure S1.**
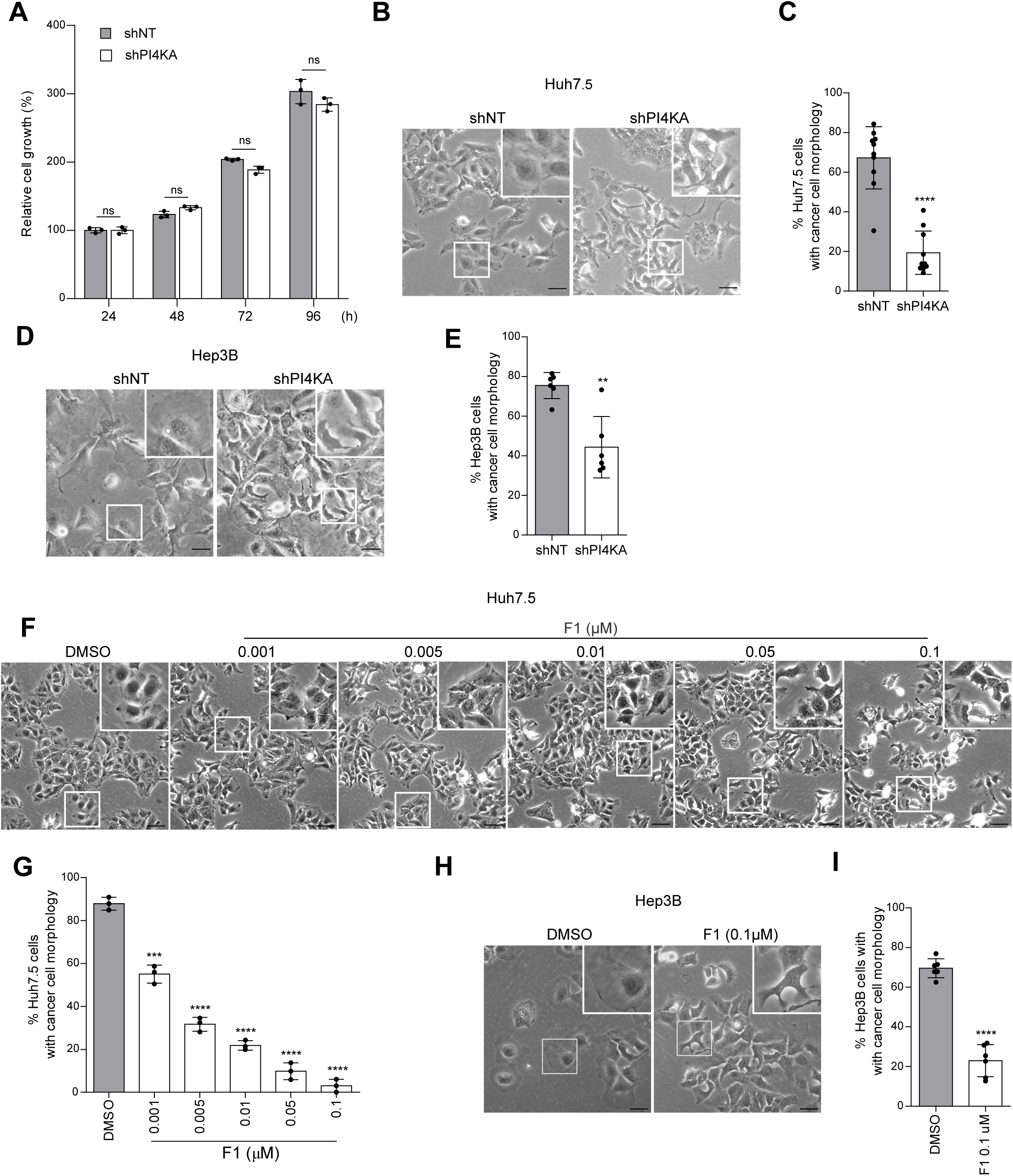
PI4KIIIα links to cell morphology. (**A**) Cell proliferation of Huh7-Lunet expressing shNT or shPI4KA. (**B-E**) Huh7.5 cells (B, C) or Hep3B cells (D, E) stably expressing shNT or shPI4KA were imaged for cell morphology (B, D) and cancer cell morphology was quantified (C, E). (**F-G**) Huh7.5 cells were either treated with DMSO or the indicated concentrations of PI4KA-F1 for 48 h. Cell morphology was recorded (F) and cancer cell morphology was assessed (G). (**H-I**) Hep 3B cells were treated with DMSO or 0.1 µM of PI4KA-F1 for 48 h and analyzed as in panels (F, G). The inserts show magnifications of the boxed areas. All scale bars represent 50 µm.

**Figure S2.**
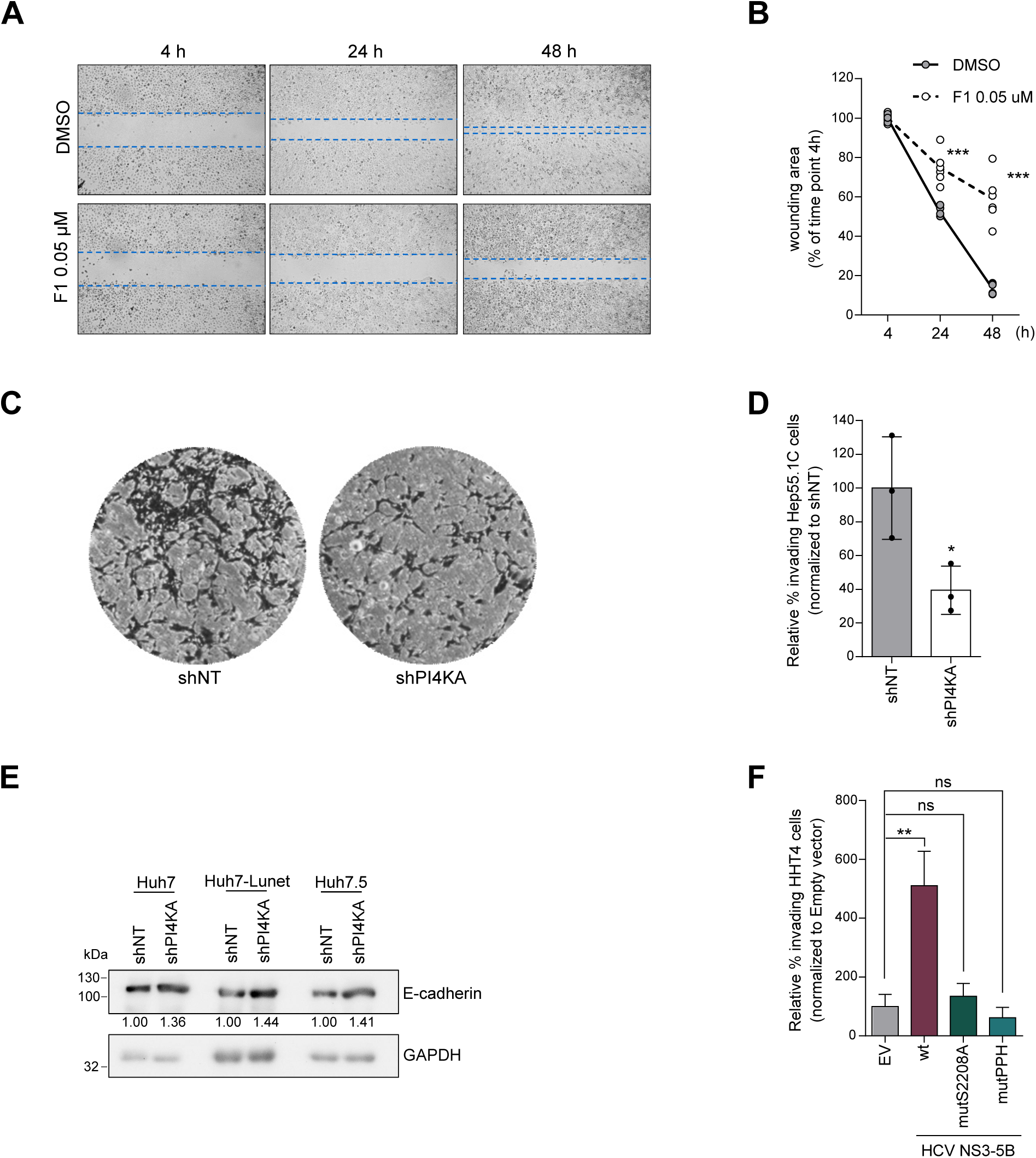
PI4KIIIα facilitates liver cancer cell motility. (**A-B**) Huh7-Lunet cells were seeded in culture-inserts 2 well and treated with DMSO or PI4KA-F1. Cell-free gaps were imaged (A) and wounding areas were quantified (B). (**C- D**) Invasion assay using Matrigel invasion chambers were performed with the murine hepatoma cell line Hep55.1C. Invading cells were imaged (C) and relative cell invasion rate was evaluated (D). (**E**) Levels of E-cadherin and GAPDH were analyzed by Western Blot in Huh7, Huh7-Lunet and Huh7.5 cells expressing shNT or shPI4KA. (**F**) Relative percentage of invasive HHT4 cells expressing the indicated HCV NS3-5B variants or empty vector.

**Figure S3.**
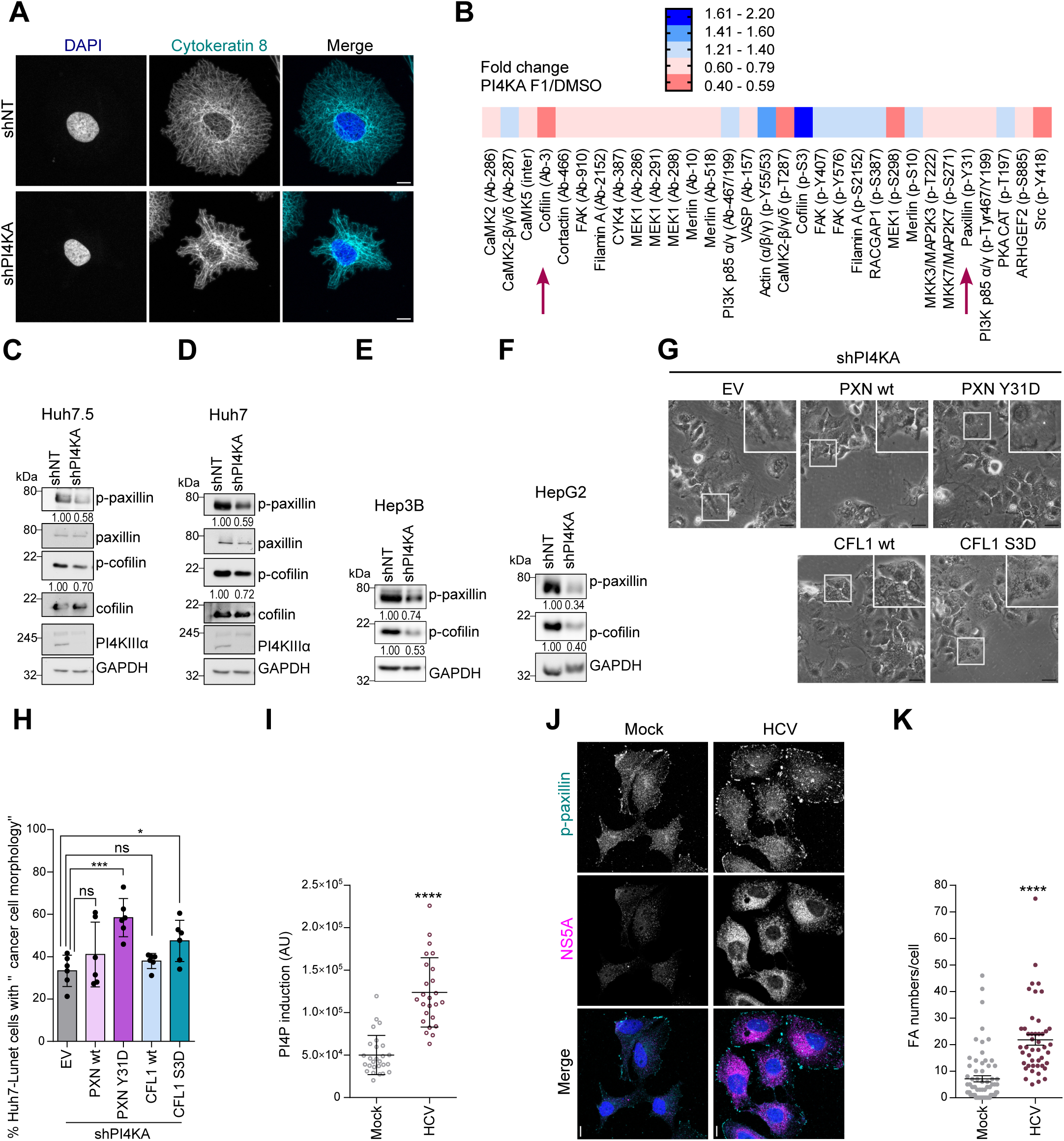
The cytoskeletal network is regulated by PI4KIIIα via an enhanced phosphorylation of paxillin and cofilin. (**A**) Huh7-Lunet-shNT and –shPI4KA cells were stained for cytokeratin-8 and DAPI and imaged by confocal laser scanning microscopy. (**B**) Huh7-Lunet cells were treated with DMSO or 0.05 µM of PI4KA-F1 for 48 h. Cell lysates were subjected to a cytoskeleton phospho antibody array (Full Moon Biosystems). Changes in expression of cytoskeletal proteins (either total or phosphorylated forms) are presented in the heatmap. (**C-F**) Levels of p-paxillin/paxillin, p-cofilin/cofilin, PI4KIIIα and GAPDH were analyzed by WB as indicated in Huh7.5 (C), Huh7 (D), Hep3B (E) and HepG2 cells (F) expressing shNT versus shPI4KA. Numbers beneath lanes represent relative quantifications of the respective band intensities. (**G-H**) Huh7-Lunet-shPI4KA cells were transduced with lentiviral vectors encoding the indicated variants of paxillin or cofilin or with empty vector (EV). Cell morphology was imaged (H) and quantified for changes in cancer cell morphology (I). Note that *PXN Y31D* and *CFL1 S3D* represent phospho mimetic mutants phenocopying the phosphorylated state of the proteins. (**I**) Huh7.5 cells were either infected with HCV or mock infected for 4 days and stained for PI4P, followed by imaging using confocal microscopy to quantify PI4P levels. (**J-K**) Huh7.5 cells, either mock or HCV-infected were stained with p-paxillin specific antibodies and phalloidin to identify FAs (J) and the FA numbers per cell were quantified in more than 40 cells (K). The inserts show magnifications of the boxed areas. Scale bars represent 10 µm in panels (A) and (I) and 50 µm in panel G.

**Figure S4.**
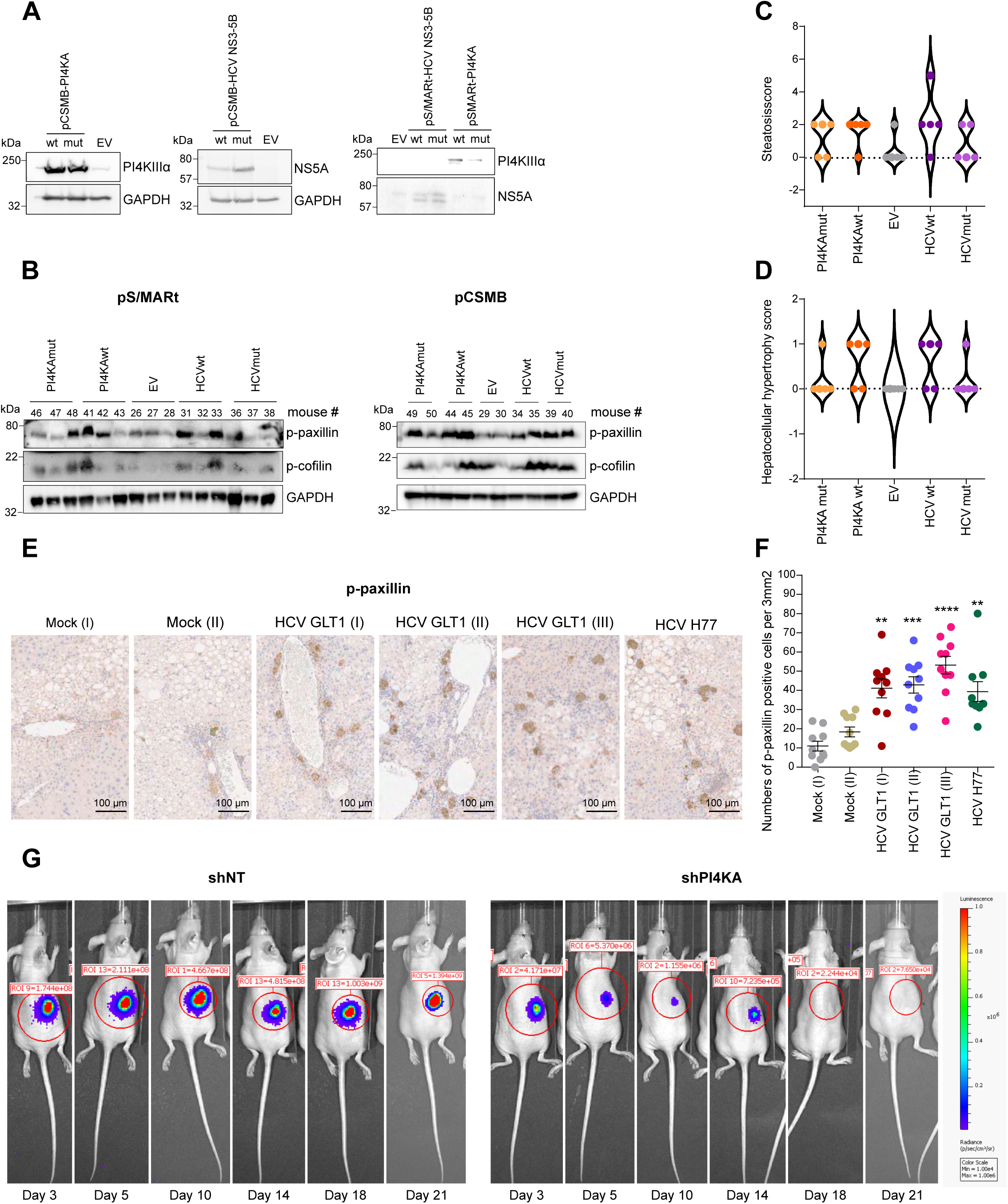
PI4KIIIα expression/activity and phosphorylation levels of paxillin and cofilin correlate *in vivo*. (**A-D**) pCSMB and pS/MARt vectors encoding wt or a mutant variant of PI4KIIIα or HCV NS3-5B or empty vector (EV) as indicated were either transfected in Huh7-Lunet cells (A) or mice were transduced by hydrodynamic tail vein injection. Cell lysates (A) or liver homogenates of transduced mice (B) were analyzed by WB using antibodies specific to the indicated proteins and mice livers were histologically evaluated using a semiquantitative scoring system (C, D). Scores for steatosis (C) and hepatocellular hypertrophy (D) are shown for each treatment group. (**E-F**) Human-liver chimeric mice were infected with HCV patient serum gt 1b, strain GLT1 or gt1a, strain mH77c. Consecutive sections from the livers were stained for p-paxillin, HCV NS5A or human albumin. Examples of p-paxillin staining in one repopulated area for each mouse are shown (E). 10 random areas from each liver were used to quantify the number of p- paxillin positive cells (F). (**G**) Huh7-Lunet-Luc-shNT and -shPI4KA cells were injected subcutaneously into the dorsal flank of nude mice. IVIS was performed at day 3, 5, 10, 14, 18 and 21. Shown are examples of one mouse for each group.

**Figure S5.**
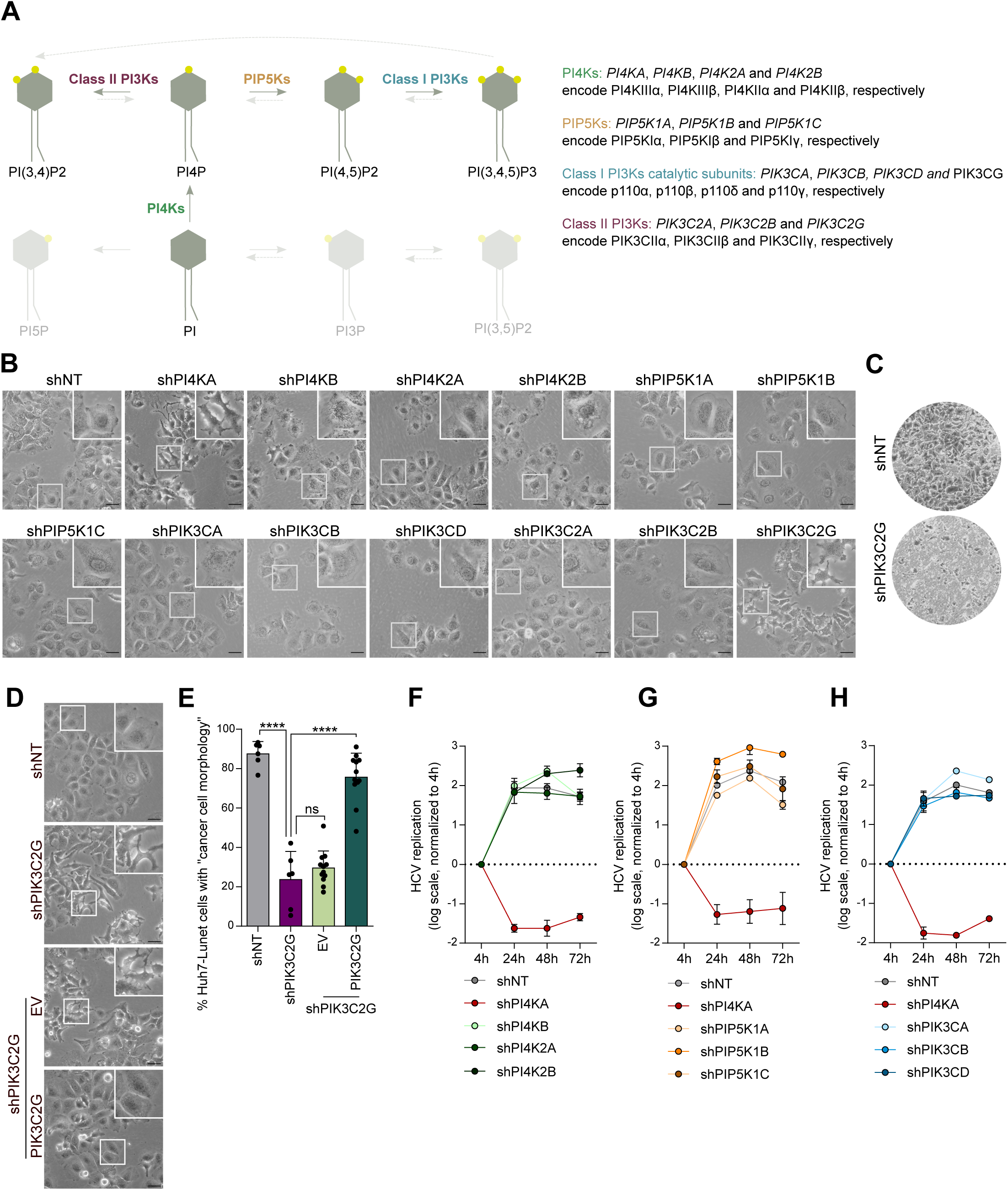
Identification of PIK3C2γ as a downstream partner of PI4KIIIα. (**A**) Schematic illustration of phosphoinositide conversion by lipid kinases (solid lines) and phosphatases (dashed lines) (**B**) Huh7-Lunet cells stably expressing the indicated shRNAs were imaged for cell morphology. (**C**) Huh7-Lunet cells expressing shNT or shPIK3C2G were analyzed for relative cell invasion rates. For quantification see Figure 5D. (**D-E**) An shRNA resistant mutant of PIK3C2G was ectopically expressed in Huh7- Lunet cells with stable knockdown of PIK3C2G. Cell morphology was analyzed (D) and cancer cell morphology was quantified (E). (**F-H**) Huh7-Lunet cells stably expressing the respective shRNAs were electroporated with HCV subgenomic reporter replicons and replication efficacy was assessed by quantification of luciferase activity at the indicated time points. The inserts show magnifications of the boxed areas. All scale bars represent 50 µm.

**Figure S6.**
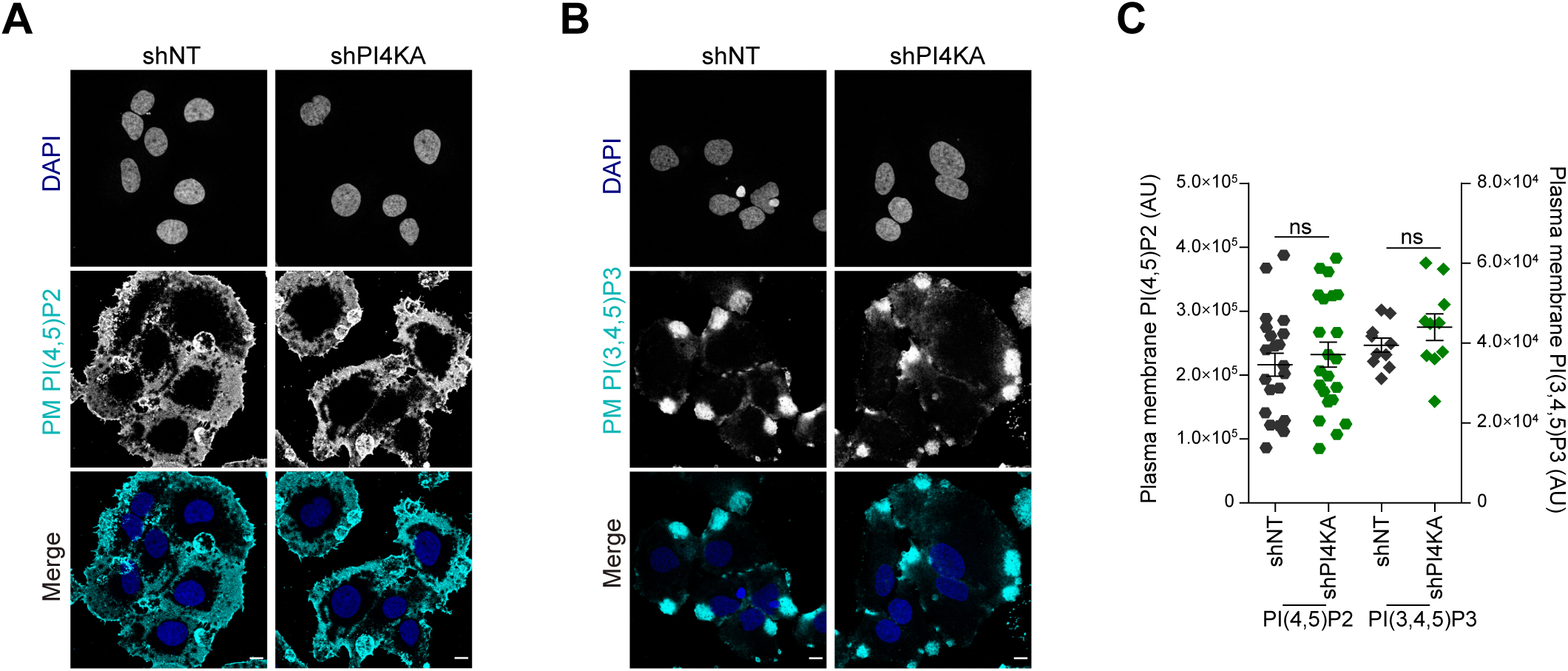
PI(3,4)P2 pools at the cell PM are dependent on PI4KIIIα expression and activity. (**A-C**) PM staining of PI(4,5)P2 (A) and PI(3,4,5)P3 (B) in Huh7-Lunet shNT versus shPI4KA and quantification of relative levels of these lipids (C). Each dot represents a single cell (n ≥ 10). All scale bars represent 10 µm.

**Figure S7.**
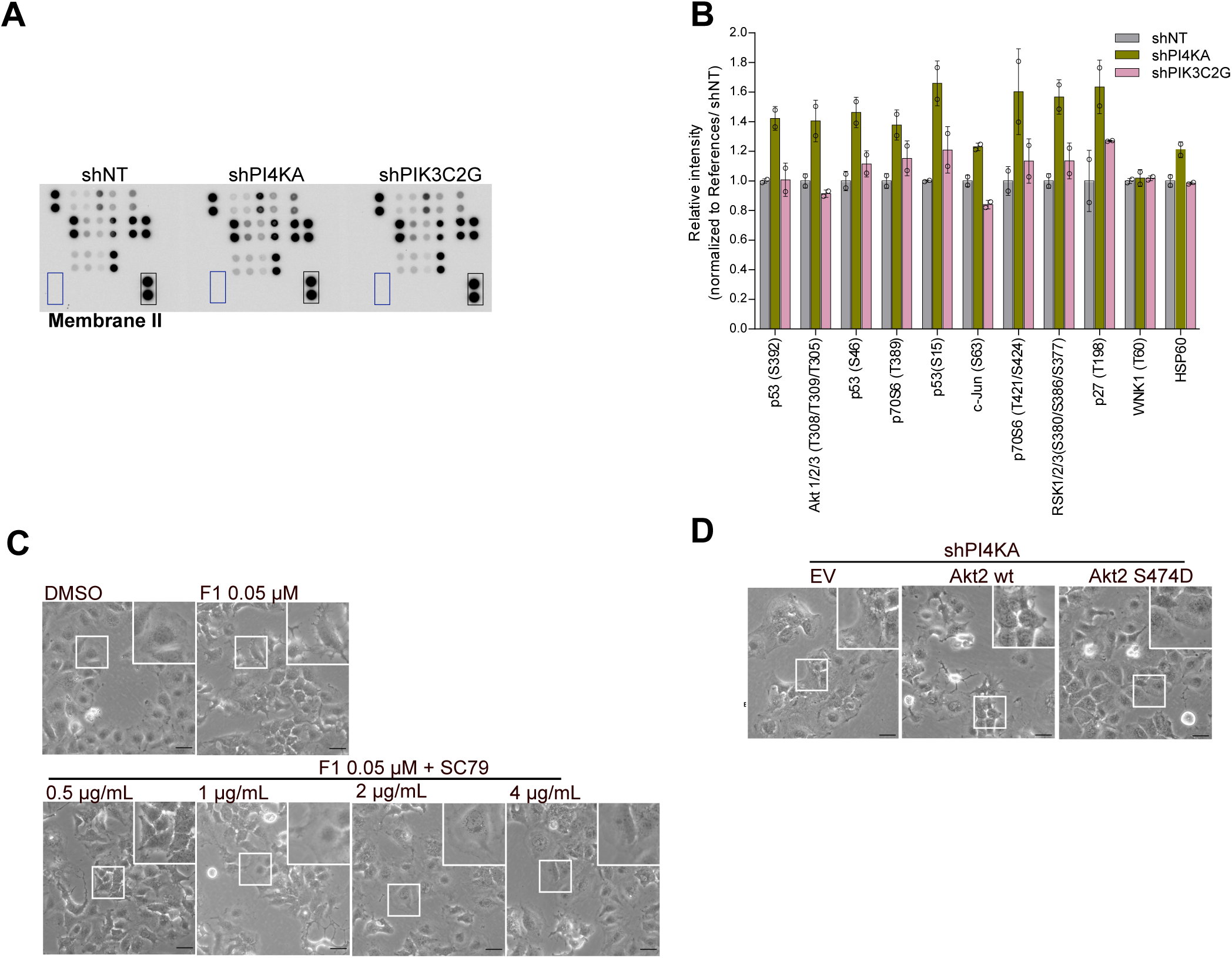
Akt2 is the mediator linking signaling from PI4KIIIα and PIK3C2γ to paxillin and further downstream. (**A and B**) The membrane II of the Proteome Profile human phospho kinase array kit was incubated with protein lysates from Huh7-Lunet expressing shNT, shPI4KA or shPIK3C2G (A). Relative signal intensities of each protein were quantified (B). (**C**) Huh7-Lunet cells were treated with PI4KA-F1 at 0.05 µM or DMSO for 48 h. Then, Akt activator SC79 at the indicated concentrations was added and cell morphology was recorded. (**D**) Akt2 variants or empty vector were expressed in Huh7-Lunet-shPI4KA using adenoviral transduction and cell morphology was recorded. The inserts show magnifications of the boxed areas. All scale bars represent 50 µm.

## Notes

### Competing Interest Statement

The authors have declared no competing interest.

## REFERENCES

[1] Rumgay H, Arnold M, Ferlay J, Lesi O, Cabasag CJ, Vignat J, et al. Global burden of primary liver cancer in 2020 and predictions to 2040. J Hepatol 2022;77:1598–1606.

[2] Hall A. The cytoskeleton and cancer. Cancer Metastasis Rev 2009;28:5–14.

[3] Yin HL, Janmey PA. Phosphoinositide regulation of the actin cytoskeleton. Annu Rev Physiol 2003;65:761–789.

[4] Bunney TD, Katan M. Phosphoinositide signalling in cancer: beyond PI3K and PTEN. Nat Rev Cancer 2010;10:342–352.

[5] D’Angelo G, Vicinanza M, Di Campli A, De Matteis MA. The multiple roles of PtdIns(4)P -- not just the precursor of PtdIns(4,5)P2. J Cell Sci 2008;121:1955–1963.

[6] Clayton EL, Minogue S, Waugh MG. Mammalian phosphatidylinositol 4-kinases as modulators of membrane trafficking and lipid signaling networks. Prog Lipid Res 2013;52:294–304.

[7] Harak C, Radujkovic D, Taveneau C, Reiss S, Klein R, Bressanelli S, et al. Mapping of functional domains of the lipid kinase phosphatidylinositol 4-kinase type III alpha involved in enzymatic activity and hepatitis C virus replication. J Virol 2014;88:9909–9926.

[8] Bojjireddy N, Botyanszki J, Hammond G, Creech D, Peterson R, Kemp DC, et al. Pharmacological and genetic targeting of the PI4KA enzyme reveals its important role in maintaining plasma membrane phosphatidylinositol 4-phosphate and phosphatidylinositol 4,5-bisphosphate levels. J Biol Chem 2014;289:6120–6132.

[9] Ilboudo A, Nault JC, Dubois-Pot-Schneider H, Corlu A, Zucman-Rossi J, Samson M, et al. Overexpression of phosphatidylinositol 4-kinase type IIIalpha is associated with undifferentiated status and poor prognosis of human hepatocellular carcinoma. BMC Cancer 2014;14:7.

[10] Harak C, Meyrath M, Romero-Brey I, Schenk C, Gondeau C, Schult P, et al. Tuning a cellular lipid kinase activity adapts hepatitis C virus to replication in cell culture. Nat Microbiol 2016;2:16247.

[11] Ziyad S, Riordan JD, Cavanaugh AM, Su T, Hernandez GE, Hilfenhaus G, et al. A Forward Genetic Screen Targeting the Endothelium Reveals a Regulatory Role for the Lipid Kinase Pi4ka in Myelo- and Erythropoiesis. Cell Rep 2018;22:1211–1224.

[12] Fuste NP, Fernandez-Hernandez R, Cemeli T, Mirantes C, Pedraza N, Rafel M, et al. Cytoplasmic cyclin D1 regulates cell invasion and metastasis through the phosphorylation of paxillin. Nat Commun 2016;7:11581.

[13] Wu DW, Wu TC, Wu JY, Cheng YW, Chen YC, Lee MC, et al. Phosphorylation of paxillin confers cisplatin resistance in non-small cell lung cancer via activating ERK- mediated Bcl-2 expression. Oncogene 2014;33:4385–4395.

[14] Liu Z, Chu S, Yao S, Li Y, Fan S, Sun X, et al. CD74 interacts with CD44 and enhances tumorigenesis and metastasis via RHOA-mediated cofilin phosphorylation in human breast cancer cells. Oncotarget 2016;7:68303–68313.

[15] Lv D, Li L, Lu Q, Li Y, Xie F, Li H, et al. PAK1-cofilin phosphorylation mediates human lung adenocarcinoma cells migration induced by apelin-13. Clin Exp Pharmacol Physiol 2016;43:569–579.

[16] Bell JB, Podetz-Pedersen KM, Aronovich EL, Belur LR, McIvor RS, Hackett PB. Preferential delivery of the Sleeping Beauty transposon system to livers of mice by hydrodynamic injection. Nat Protoc 2007;2:3153–3165.

[17] Argyros O, Wong SP, Niceta M, Waddington SN, Howe SJ, Coutelle C, et al. Persistent episomal transgene expression in liver following delivery of a scaffold/matrix attachment region containing non-viral vector. Gene Ther 2008;15:1593–1605.

[18] Meuleman P, Libbrecht L, De Vos R, de Hemptinne B, Gevaert K, Vandekerckhove J, et al. Morphological and biochemical characterization of a human liver in a uPA-SCID mouse chimera. Hepatology 2005;41:847–856.

[19] Burke JE. Structural Basis for Regulation of Phosphoinositide Kinases and Their Involvement in Human Disease. Mol Cell 2018;71:653–673.

[20] Hammond GR, Schiavo G, Irvine RF. Immunocytochemical techniques reveal multiple, distinct cellular pools of PtdIns4P and PtdIns(4,5)P(2). Biochem J 2009;422:23–35.

[21] Feng Z, Yu CH. PI(3,4)P(2)-mediated membrane tubulation promotes integrin trafficking and invasive cell migration. Proc Natl Acad Sci U S A 2021;118.

[22] Lopez-Colome AM, Lee-Rivera I, Benavides-Hidalgo R, Lopez E. Paxillin: a crossroad in pathological cell migration. J Hematol Oncol 2017;10:50.

[23] Liu Z, Tian Y, Machida K, Lai MM, Luo G, Foung SK, et al. Transient activation of the PI3K-AKT pathway by hepatitis C virus to enhance viral entry. J Biol Chem 2012;287:41922–41930.

[24] Mannova P, Beretta L. Activation of the N-Ras-PI3K-Akt-mTOR pathway by hepatitis C virus: control of cell survival and viral replication. J Virol 2005;79:8742–8749.

[25] Shi Q, Hoffman B, Liu Q. PI3K-Akt signaling pathway upregulates hepatitis C virus RNA translation through the activation of SREBPs. Virology 2016;490:99–108.

[26] Guo P, He Y, Chen L, Qi L, Liu D, Chen Z, et al. Cytosolic phospholipase A2alpha modulates cell-matrix adhesion via the FAK/paxillin pathway in hepatocellular carcinoma. Cancer Biol Med 2019;16:377–390.

[27] Zhang L, Chai Z, Kong S, Feng J, Wu M, Tan J, et al. Nujiangexanthone A Inhibits Hepatocellular Carcinoma Metastasis via Down Regulation of Cofilin 1. Front Cell Dev Biol 2021;9:644716.

[28] Hawkins PT, Stephens LR. Emerging evidence of signalling roles for PI(3,4)P2 in Class I and II PI3K-regulated pathways. Biochem Soc Trans 2016;44:307–314.

[29] Hasegawa J, Tokuda E, Tenno T, Tsujita K, Sawai H, Hiroaki H, et al. SH3YL1 regulates dorsal ruffle formation by a novel phosphoinositide-binding domain. J Cell Biol 2011;193:901–916.

[30] Fukumoto M, Ijuin T, Takenawa T. PI(3,4)P(2) plays critical roles in the regulation of focal adhesion dynamics of MDA-MB-231 breast cancer cells. Cancer Sci 2017;108:941–951.

[31] Braccini L, Ciraolo E, Campa CC, Perino A, Longo DL, Tibolla G, et al. PI3K- C2gamma is a Rab5 effector selectively controlling endosomal Akt2 activation downstream of insulin signalling. Nat Commun 2015;6:7400.

[32] Ono F, Nakagawa T, Saito S, Owada Y, Sakagami H, Goto K, et al. A novel class II phosphoinositide 3-kinase predominantly expressed in the liver and its enhanced expression during liver regeneration. J Biol Chem 1998;273:7731–7736.

[33] De Santis MC, Gozzelino L, Margaria JP, Costamagna A, Ratto E, Gulluni F, et al. Lysosomal lipid switch sensitises to nutrient deprivation and mTOR targeting in pancreatic cancer. Gut 2023;72:360–371.

[34] Thibault B, Ramos-Delgado F, Guillermet-Guibert J. Targeting Class I-II-III PI3Ks in Cancer Therapy: Recent Advances in Tumor Biology and Preclinical Research. Cancers (Basel) 2023;15.

[35] Ruzzenente A, Fassan M, Conci S, Simbolo M, Lawlor RT, Pedrazzani C, et al. Cholangiocarcinoma Heterogeneity Revealed by Multigene Mutational Profiling: Clinical and Prognostic Relevance in Surgically Resected Patients. Ann Surg Oncol 2016;23:1699–1707.

[36] Kim JH, Hwang KH, Eom M, Kim M, Park EY, Jeong Y, et al. WNK1 promotes renal tumor progression by activating TRPC6-NFAT pathway. FASEB J 2019;33:8588–8599.

[37] Sbrissa D, Semaan L, Govindarajan B, Li Y, Caruthers NJ, Stemmer PM, et al. A novel cross-talk between CXCR4 and PI4KIIIalpha in prostate cancer cells. Oncogene 2019;38:332–344.

[38] Ishikawa S, Egami H, Kurizaki T, Akagi J, Tamori Y, Yoshida N, et al. Identification of genes related to invasion and metastasis in pancreatic cancer by cDNA representational difference analysis. J Exp Clin Cancer Res 2003;22:299–306.

[39] Wang C, Che L, Hu J, Zhang S, Jiang L, Latte G, et al. Activated mutant forms of PIK3CA cooperate with RasV12 or c-Met to induce liver tumour formation in mice via AKT2/mTORC1 cascade. Liver Int 2016;36:1176–1186.

[40] Fayard E, Xue G, Parcellier A, Bozulic L, Hemmings BA. Protein kinase B (PKB/Akt), a key mediator of the PI3K signaling pathway. Curr Top Microbiol Immunol 2010;346:31–56.

[41] Nitulescu GM, Margina D, Juzenas P, Peng Q, Olaru OT, Saloustros E, et al. Akt inhibitors in cancer treatment: The long journey from drug discovery to clinical use (Review). Int J Oncol 2016;48:869–885.

[42] Garofalo RS, Orena SJ, Rafidi K, Torchia AJ, Stock JL, Hildebrandt AL, et al. Severe diabetes, age-dependent loss of adipose tissue, and mild growth deficiency in mice lacking Akt2/PKB beta. J Clin Invest 2003;112:197–208.

[43] Park Y, Park JM, Kim DH, Kwon J, Kim IA. Inhibition of PI4K IIIalpha radiosensitizes in human tumor xenograft and immune-competent syngeneic murine tumor model. Oncotarget 2017;8:110392–110405.

[44] Freitag A, Prajwal P, Shymanets A, Harteneck C, Nurnberg B, Schachtele C, et al. Development of first lead structures for phosphoinositide 3-kinase-C2gamma inhibitors. J Med Chem 2015;58:212–221.

[45] Khera L, Paul C, Kaul R. Hepatitis C Virus Mediated Metastasis in Hepatocellular Carcinoma as a Therapeutic Target for Cancer Management. Curr Drug Metab 2018;19:224–235.

[46] Reiss S, Rebhan I, Backes P, Romero-Brey I, Erfle H, Matula P, et al. Recruitment and activation of a lipid kinase by hepatitis C virus NS5A is essential for integrity of the membranous replication compartment. Cell Host Microbe 2011;9:32–45.

[47] Harouaka D, Engle RE, Wollenberg K, Diaz G, Tice AB, Zamboni F, et al. Diminished viral replication and compartmentalization of hepatitis C virus in hepatocellular carcinoma tissue. Proc Natl Acad Sci U S A 2016;113:1375–1380.

[48] Goossens N, Hoshida Y. Hepatitis C virus-induced hepatocellular carcinoma. Clin Mol Hepatol 2015;21:105–114.

## SUPPLEMENTAL REFERENCES

[1] Kleine M, Riemer M, Krech T, DeTemple D, Jager MD, Lehner F, et al. Explanted diseased livers - a possible source of metabolic competent primary human hepatocytes. PLoS One 2014;9:e101386.

[2] Reiss S, Rebhan I, Backes P, Romero-Brey I, Erfle H, Matula P, et al. Recruitment and activation of a lipid kinase by hepatitis C virus NS5A is essential for integrity of the membranous replication compartment. Cell Host Microbe 2011;9:32–45.

[3] Reiss S, Harak C, Romero-Brey I, Radujkovic D, Klein R, Ruggieri A, et al. The lipid kinase phosphatidylinositol-4 kinase III alpha regulates the phosphorylation status of hepatitis C virus NS5A. PLoS Pathog 2013;9:e1003359.

[4] Harak C, Meyrath M, Romero-Brey I, Schenk C, Gondeau C, Schult P, et al. Tuning a cellular lipid kinase activity adapts hepatitis C virus to replication in cell culture. Nat Microbiol 2016;2:16247.

[5] Kato T, Date T, Murayama A, Morikawa K, Akazawa D, Wakita T. Cell culture and infection system for hepatitis C virus. Nat Protoc 2006;1:2334–2339.

[6] Lindenbach BD. Measuring HCV infectivity produced in cell culture and in vivo. Methods Mol Biol 2009;510:329–336.

[7] Hammond GR, Schiavo G, Irvine RF. Immunocytochemical techniques reveal multiple, distinct cellular pools of PtdIns4P and PtdIns(4,5)P(2). Biochem J 2009;422:23–35.

[8] Berg S, Kutra D, Kroeger T, Straehle CN, Kausler BX, Haubold C, et al. ilastik: interactive machine learning for (bio)image analysis. Nat Methods 2019;16:1226–1232.

[9] Kleiner DE, Brunt EM, Van Natta M, Behling C, Contos MJ, Cummings OW, et al. Design and validation of a histological scoring system for nonalcoholic fatty liver disease. Hepatology 2005;41:1313–1321.

[10] Meuleman P, Libbrecht L, De Vos R, de Hemptinne B, Gevaert K, Vandekerckhove J, et al. Morphological and biochemical characterization of a human liver in a uPA-SCID mouse chimera. Hepatology 2005;41:847–856.

[11] Heuss C, Rothhaar P, Burm R, Lee JY, Ralfs P, Haselmann U, et al. A Hepatitis C virus genotype 1b post-transplant isolate with high replication efficiency in cell culture and its adaptation to infectious virus production in vitro and in vivo. PLoS Pathog 2022;18:e1010472.

[12] Meuleman P, Bukh J, Verhoye L, Farhoudi A, Vanwolleghem T, Wang RY, et al. In vivo evaluation of the cross-genotype neutralizing activity of polyclonal antibodies against hepatitis C virus. Hepatology 2011;53:755–762.

[13] Graham FL, Smiley J, Russell WC, Nairn R. Characteristics of a human cell line transformed by DNA from human adenovirus type 5. J Gen Virol 1977;36:59–74.

[14] Nakabayashi H, Taketa K, Miyano K, Yamane T, Sato J. Growth of human hepatoma cells lines with differentiated functions in chemically defined medium. Cancer Res 1982;42:3858–3863.

[15] Backes P, Quinkert D, Reiss S, Binder M, Zayas M, Rescher U, et al. Role of annexin A2 in the production of infectious hepatitis C virus particles. J Virol 2010;84:5775–5789.

[16] Blight KJ, McKeating JA, Rice CM. Highly permissive cell lines for subgenomic and genomic hepatitis C virus RNA replication. J Virol 2002;76:13001–13014.

[17] Aden DP, Fogel A, Plotkin S, Damjanov I, Knowles BB. Controlled synthesis of HBsAg in a differentiated human liver carcinoma-derived cell line. Nature 1979;282:615–616.

[18] Knowles BB, Howe CC, Aden DP. Human hepatocellular carcinoma cell lines secrete the major plasma proteins and hepatitis B surface antigen. Science 1980;209:497–499.

[19] Kress S, Konig J, Schweizer J, Lohrke H, Bauer-Hofmann R, Schwarz M. p53 mutations are absent from carcinogen-induced mouse liver tumors but occur in cell lines established from these tumors. Mol Carcinog 1992;6:148–158.

